# RCD1 Coordinates Chloroplastic and Mitochondrial Electron Transfer through Interaction with ANAC Transcription Factors in Arabidopsis

**DOI:** 10.1101/327411

**Authors:** Alexey Shapiguzov, Julia P. Vainonen, Kerri Hunter, Helena Tossavainen, Arjun Tiwari, Sari Järvi, Maarit Hellman, Brecht Wybouw, Fayezeh Aarabi, Saleh Alseekh, Nina Sipari, Lauri Nikkanen, Katrien Van Der Kelen, Julia Krasensky-Wrzaczek, Jarkko Salojärvi, Mikael Brosché, Markku Keinänen, Esa Tyystjärvi, Eevi Rintamäki, Bert De Rybel, Alisdair R. Fernie, Frank Van Breusegem, Perttu Permi, Eva-Mari Aro, Michael Wrzaczek, Jaakko Kangasjärvi

## Abstract

Signaling from chloroplasts and mitochondria, both dependent on reactive oxygen species (ROS), merge at the nuclear protein RADICAL-INDUCED CELL DEATH1 (RCD1). ROS produced in the chloroplasts affect the abundance, thiol redox state and oligomerization of RCD1. RCD1 directly interacts *in vivo* with ANAC013 and ANAC017 transcription factors, which are the mediators of the ROS-related mitochondrial complex III retrograde signa and suppresses activity of ANAC013 and ANAC017. Inactivation of *RCD1* leads to increased expression of ANAC013 and ANAC017-regulated genes belonging to the mitochondrial dysfunction stimulon (MDS), including genes for mitochondrial alternative oxidases *(AOXs).* Accumulating AOXs and other MDS gene products alter electron transfer pathways in the chloroplasts, leading to diminished production of chloroplastic ROS and increased protection of photosynthetic apparatus from ROS damage. RCD1-dependent regulation affects chloroplastic and mitochondrial retrograde signaling including chloroplast signaling by 3’-phosphoadenosine 5’-phosphate (PAP). Sensitivity of RCD1 to organellar ROS provides feedback control of nuclear gene expression.

## Introduction

Cells of photosynthesizing eukaryotes are unique in harboring two types of energy organelles, chloroplasts and mitochondria, which are complexly intertwined. On the one hand, they interact at an operational level by the exchange of metabolites, energy and reducing power (Noguchi and Yoshida 2008, Cardol et al., 2009, Bailleul et al., 2015). Reducing power flows between the organelles through several pathways including photorespiration (Watanabe et al., 2016), malate shuttles (Scheibe 2004, Zhao et al., 2018) as well as export of carbohydrates from the chloroplasts and import into the mitochondria. Furthermore, mitochondrial alternative oxidases (AOXs) are known to consume the excess reducing power generated in chloroplasts during photosynthesis, thereby both acting as safety valves and facilitating optimal rates of photosynthesis (Yoshida et al., 2006, Vanlerberghe et al., 2016). On the other hand, chloroplasts and mitochondria also interact at the signaling level. The so-called retrograde signaling pathways originate from the organelles and influence the expression of nuclear genes (de Souza et al., 2016, Leister 2017, Waszczak et al., 2018). These pathways provide feedback communication between the organelles and the gene expression apparatus in the nucleus to adjust expression of genes encoding organelle components in accordance with demands arising from developmental stage or environmental conditions.

One class of compounds playing pivotal roles in plant organellar signaling both from chloroplasts and mitochondria are reactive oxygen species (ROS) (Dietz et al., 2016, Noctor et al., 2017, Waszczak et al. 2018). ROS are inevitable by-products of aerobic energy metabolism. Superoxide anion radical (O_2_^▪-^) is formed in the organelles by the occasional leakage of electrons from the organellar electron transfer chains (ETCs) to molecular oxygen (O_2_). For instance, in illuminated chloroplasts reduction of O_2_ to superoxide anion by Photosystem I (PSI) initiates the water-water cycle (Asada 2006, Awad et al., 2015). In this process, O_2_^▪-^ is enzymatically converted to hydrogen peroxide (H_2_O_2_) which is subsequently reduced to water by chloroplastic H_2_O_2_-scavenging systems. One of the widely employed experimental means to boost chloroplast production of ROS is to apply methyl viologen (MV), a chemical that catalyzes shuttling of electrons from PSI to O_2_ (Farrington et al., 1973). The immediate product of this reaction, the superoxide anion, is not likely to directly mediate organellar signaling; however, H_2_O_2_ is involved in many retrograde signaling pathways (Leister 2017, Waszczak et al. 2018). Organellar H_2_O_2_ can be translocated to the nucleus directly (Caplan et al., 2015, Exposito-Rodriguez et al., 2017) or it can oxidize thiol groups of specific proteins, thereby converting the ROS signal into thiol redox signals (Møller and Kristensen 2004, Nietzel et al., 2017). One recently discovered process targeted by chloroplastic H_2_O_2_ is the metabolism of 3’-phosphoadenosine 5’-phosphate (PAP). PAP is a toxic by-product of sulfate metabolism produced when cytoplasmic sulfotransferases (SOTs) transfer a sulfuryl group from PAP-sulfate (PAPS) to various target compounds (Klein and Papenbrock 2004). PAP is transported to chloroplasts where it is detoxified by dephosphorylation to adenosine monophosphate in a reaction catalyzed by the adenosine bisphosphate phosphatase 1, SAL1 (Quintero et al., 1996, Chan et al., 2016). Oxidation of SAL1 thiols by chloroplastic H_2_O_2_ inactivates the enzyme and accumulating PAP thereby acts as a retrograde signal (Chan et al. 2016). How PAP initiates transcriptional reprogramming remains unclear, however it has been suggested that inhibition of the activity of nuclear RNA 5’-3’ exonucleases is involved (Estavillo et al., 2011).

ROS are also produced in the mitochondria, for example by complex III at the outer side of the inner mitochondrial membrane (Cvetkovska et al., 2013, Ng et al., 2014, Huang et al., 2016). Such ROS are implicated in mitochondrial retrograde signaling. Blocking electron transfer through complex III *via* application of the inhibitors antimycin A (AA) or myxothiazol (myx) enhances electron leakage and thus the mitochondrial retrograde signal. Two known mediators of this signal are the transcription factors ANAC013 (De Clercq et al., 2013) and ANAC017 (Ng et al., 2013, Van Aken et al., 2016) that are both bound to the endoplasmic reticulum (ER) by their transmembrane domain. Mitochondria-derived signals lead to proteolytic cleavage of this domain, such that the proteins are released from the ER, and then translocated to the nucleus where they activate the mitochondrial dysfunction stimulon (MDS) genes, including AOX. Accumulation of AOX likely enhances the activity of the alternative respiratory pathway, which is able to bypass complex III. The *ANAC013* gene is also itself an MDS gene, generating a positive feedback loop within the signaling pathway (De Clercq et al. 2013, Van Aken et al., 2016).

Whereas multiple retrograde signaling pathways have been described in detail (de Souza et al. 2016, Leister 2017, Waszczak et al. 2018), it is still largely unknown how the numerous chloroplast- and mitochondria-derived signals are cooperatively processed by the nuclear gene expression system. Nuclear cyclin-dependent kinase E is implicated in the expression of both chloroplastic *(LHCB2.4)* and mitochondrial *(AOX1a)* components in response to perturbations of chloroplast ETC (Blanco et al., 2014), mitochondrial ETC, or H_2_O_2_ treatment (Ng et al., 2013). The transcription factor ABI4 responds to retrograde signals from both of these organelles (Giraud et al., 2009, Blanco et al. 2014). Mitochondrial signaling *via* ANAC017 was recently suggested to converge with chloroplast PAP signaling, based on similarities in their transcriptomic profiles (Van Aken and Pogson 2017); the mechanistic details underlying this convergence remain currently unknown.

Arabidopsis RADICAL-INDUCED CELL DEATH1 (RCD1) is a nuclear protein containing a WWE, a PARP-like [poly (ADP-ribose) polymerase-like], and a C-terminal RST domain (<u>R</u>CD1-<u>S</u>RO1-<u>T</u>AF4) (Jaspers et al., 2009, Jaspers et al., 2010). In yeast two-hybrid studies RCD1 interacted with many transcription factors including ANAC013, DREB2A (Vainonen et al., 2012), Rap2.4a (Hiltscher et al., 2014) and others (Jaspers et al. 2009) *via* the RST domain (Jaspers et al., 2010). In agreement with the multiple putative interaction partners of RCD1, the *rcd1* mutant demonstrates pleiotropic phenotypes in diverse stress and developmental responses (Jaspers et al. 2009). It has been identified in screens for sensitivity to ozone (Overmyer et al., 2000), tolerance to MV (Fujibe et al., 2004) and redox imbalance in the chloroplasts (Heiber et al., 2007, Hiltscher et al. 2014). Under standard growth conditions, the *rcd1* mutant displays differential expression of over 400 genes, including those encoding mitochondrial AOXs (Jaspers et al. 2009, Brosché et al., 2014) and the chloroplast 2-Cys peroxiredoxin (2-CP) (Heiber et al. 2007, Hiltscher et al. 2014).

Here we have addressed the role of RCD1 in integration of ROS signals emitted by both mitochondria and chloroplasts. Abundance, redox status and oligomerization state of the nuclear-localized RCD1 protein changed in response to ROS generated in the chloroplasts. Furthermore, RCD1 directly interacted *in vivo* with ANAC013 and ANAC017 functioning as a negative regulator of both transcription factors. We demonstrate that RCD1 is a molecular component which integrates organellar signal input from both chloroplasts and mitochondria to exert its influence on nuclear gene expression.

## Results

### The response to chloroplastic ROS is compromised in *rcd1*

In illuminated chloroplasts Photosystem I (PSI) is the main producer of ROS. Methyl viologen (MV) redirects electrons from PSI to form ROS, which triggers a chain of reactions that ultimately lead to inhibition of Photosystem II (PSII) (Farrington et al. 1973, Nishiyama et al., 2011). To reveal the significance of RCD1 in responses to PSI-produced chloroplastic ROS, rosettes of Arabidopsis were pre-treated overnight in darkness with MV and exposed to light. After the dark period, the plants displayed unchanged PSII photochemical yield (Fv/Fm). Subsequent exposure to three hours of light resulted in decrease of Fv/Fm in wild type (Col-0) (Fig. 1A), but not in the *rcd1* mutant. Moreover, tolerance of *rcd1* was evident under various concentrations of MV (Fig. 1B). The superoxide anion is an unstable compound that is enzymatically reduced to the more long-lived H_2_O_2_. Chloroplastic production of H_2_O_2_ in the presence of MV in the light was assessed by staining plants with 3,3’-diaminobenzidine (DAB). After pre-treatment with MV, higher H_2_O_2_ accumulation was evident in both Col-0 and *rcd1* (Fig. S1A). Subsequent incubation of these plants under light led to a time-dependent increase in the H_2_O_2_ accumulation in Col-0, but not in *rcd1.*

**Figure 1.**
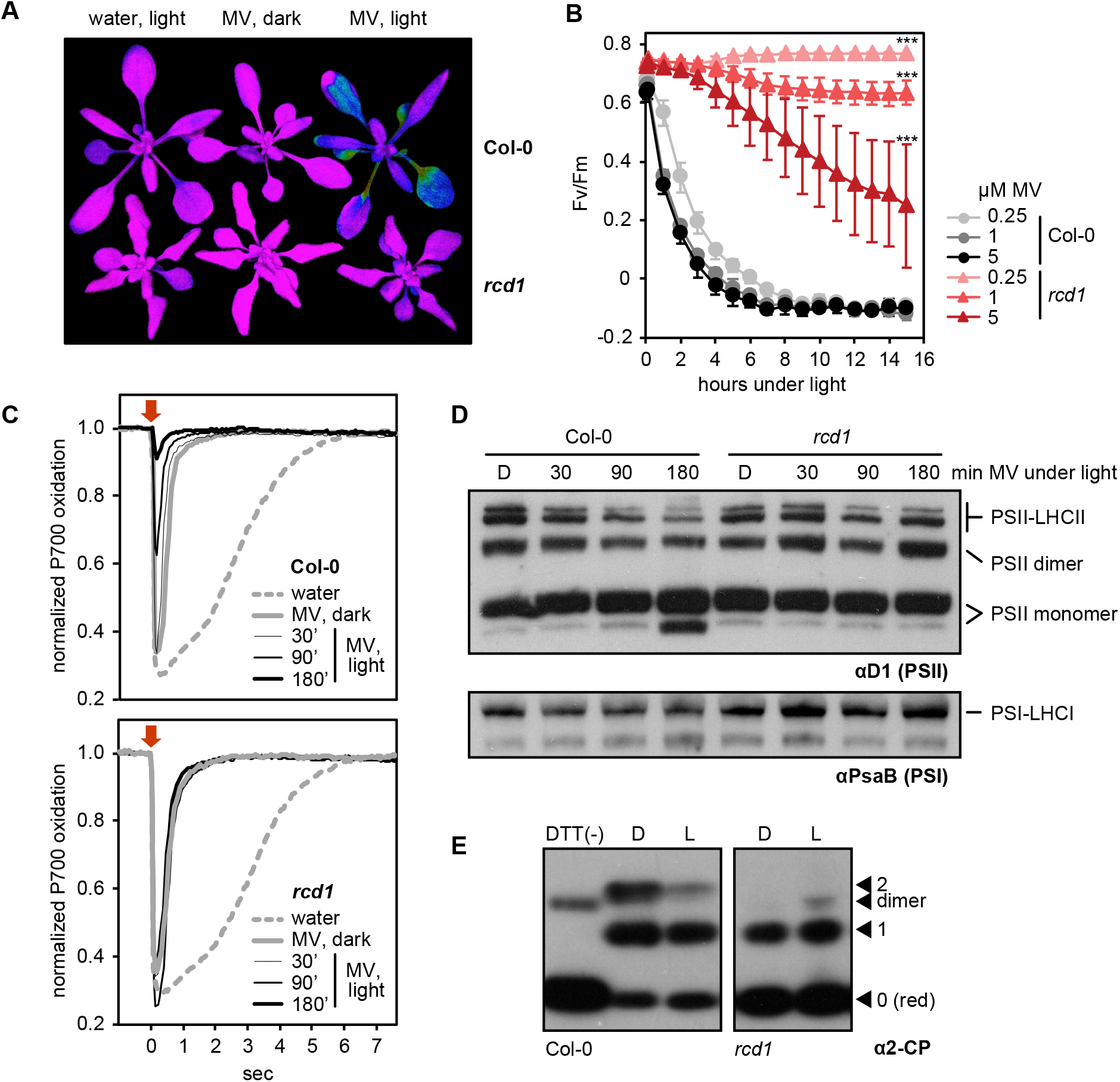
Changed tolerance to ROS of photosynthetic apparatus in *rcd1.* (A) Effect of MV (1 μM here and elsewhere unless stated otherwise) on photochemical yield of PSII in the darkness and under light, as seen in false-color image of Fv/Fm. (B) Tolerance of *rcd1* mutant to PSII photoinhibition by ROS under a range of MV concentrations [mean ± SD, asterisks denote values significantly different from those in the similarly treated wild type at the last time point of the assay (P value < 0.001, bonferroni post hoc correction, for full statistical analysis see Supplementary Table 5)]. (C) Effect of MV on PSI (P700) oxidation in Col-0 and *rcd1.* Flashes of PSII-specific light are labeled with arrows. (D) Immunostaining for PSII (αD1 antibody) and PSI (αPsaB antibody) in photosynthetic complexes isolated from MV-treated leaves that were exposed to light for indicated times (D – dark control). (E) Thiol bond-specific labeling of the chloroplast enzyme 2-CP in Col-0 (left) and *rcd1* (right). Proteins were isolated from leaves incubated in darkness (D), or under light (L). DTT (-) control contains predominantly unlabeled form. Unlabeled reduced (red), singly and doubly labeled oxidized forms and the putative dimer are marked.

In several MV-tolerant mutants the resistance is based on restricted access of MV to chloroplasts (Hawkes 2014). However, in *rcd1* pretreatment with MV led to initial increase in H_2_O_2_ production similar to that in the wild type (Fig. S1A), suggesting that resistance of *rcd1* was not due to restricted delivery of MV to PSI. In order to test this directly, oxidation of PSI was assessed by *in vivo* spectroscopy using DUAL-PAM. Leaves were adapted to far-red light, which is more efficiently used by PSI than PSII. Under these conditions PSI is producing electrons at a faster rate than it is supplied by electrons coming from PSII, and hence the PSI reaction center P700 becomes oxidized. Then a flash of orange light was provided that is efficiently absorbed by PSII. Electrons generated by PSII transiently reduced PSI, after which the kinetics of PSI re-oxidation was followed (Fig. 1C). Pre-treatment of leaves with MV led to a more rapid oxidation of PSI after the flash of orange light. The effect of MV was identical in Col-0 and *rcd1,* indicating unaltered access of MV to PSI in the *rcd1* mutant. Thus, the tolerance of *rcd1* likely arose from alterations taking place downstream from PSI. Indeed, in Col-0 exposure to light in the presence of MV led to a progressive decrease in the reduction of PSI resulting from the flash of orange light, suggesting a deterioration in PSII function. However, such behavior was not observed in *rcd1* (Fig. 1C).

To study this phenomenon in more detail, Col-0 and *rcd1* protein extracts were separated by SDS-PAGE followed by immunoblotting with antibodies against the PSII subunit D1 and the PSI subunit PsaB. No significant differences in PSI nor PSII abundances were detected (Fig. S1B). When the photosynthetic complexes were separated by native PAGE, immunoblotting with αD1 antibody revealed PSII species of diverse molecular weights (Fig. 1D). The largest of them corresponded to PSII associated with its light-harvesting antennae complex (LHCII) while the smallest corresponded to PSII monomers. In accordance with the results obtained with DUAL-PAM, chloroplastic ROS production in Col-0 leaves pre-treated with MV led to progressive disassembly of PSII-LHCII complexes and accumulation of PSII monomers, while no changes were observed in *rcd1* (Fig. 1D). Importantly, no signs of PSI inhibition were evident either in DUAL-PAM (Fig. 1C), or in PSI immunoblotting assays (Fig. 1D) in either genotype. The fact that inhibition of the photosynthetic apparatus by ROS manifested itself in disassembly of PSII complexes, but did not affect PSI where ROS were formed, suggested that inhibition resulted from a regulated mechanism rather than uncontrolled oxidation by ROS, and that this mechanism requires the activity of RCD1.

Previous studies have described *rcd1* as a mutant with altered ROS metabolism and redox status of the chloroplasts, although the underlying mechanisms of these defects could not be unequivocally determined (Fujibe et al. 2004, Heiber et al. 2007, Hiltscher et al. 2014). Analyses of abundance and redox states of the low molecular weight antioxidant compounds ascorbate and glutathione could not convincingly explain resistance of *rcd1* to chloroplastic ROS (Heiber et al. 2007, Hiltscher et al. 2014). In further search for possible mechanisms of MV tolerance of *rcd1,* thiol redox states of chloroplast enzymes involved in redox signaling and antioxidant defense were next assessed. 2-Cys peroxiredoxin (2-CP) is an abundant ROS-processing enzyme localized in the chloroplast stroma (Konig et al., 2002, Peltier et al., 2006, Liebthal et al., 2018). The abundance of the 2-CP protein was unchanged in *rcd1* (Fig. S1C). However, when total protein extracts were subjected to thiol bond-specific labeling, separated by SDS-PAGE and immunoblotted with an antibody against 2-CP, different redox forms of 2-CP were revealed in the studied lines (Fig. 1E). In Col-0 a fraction of 2-CP was present as a doubly oxidized form both in the darkness and under light. By contrast, in *rcd1* the doubly oxidized form of 2-CP was not detected under the tested conditions, while the unlabeled form corresponding to reduced 2-CP accumulated to higher amounts. This indicated that in *rcd1* the thiol redox state of 2-CP was, even under standard growth conditions, more reduced than in Col-0, and thus suggests that RCD1 is involved in the regulation of the chloroplastic redox status.

To understand the molecular basis of the RCD1-dependent redox alterations, the levels of proteins involved in photosynthesis and ROS scavenging were analyzed by immunoblotting. These included PSBS (non-photochemical quenching), PTOX (terminal oxidation of plastoquinone pool), PGR5, PGRL1, NDH45, NDH48, and PQL2 (thylakoid proton gradient and cyclic electron transfer). ROS-scavenging enzymes including chloroplastic and cytosolic isoforms of superoxide dismutase (SOD) and peroxiredoxin Q (PRXQ) were also tested. None of these proteins showed altered abundance in *rcd1* compared to Col-0 (Fig. S1D). The nucleotide redox couples NAD+/NADH and NADP+/NADPH play key roles in cellular redox balance and energy metabolism. To account for possible metabolic distortions in the *rcd1* mutant, NAD+, NADP+, NADH and NADPH were extracted from light-adapted Col-0 and *rcd1* rosettes, and their abundance was measured. However, no changes were detected in *rcd1* compared to the wild type (Fig. S1E). To reveal possible alterations in the distribution of these compounds between the cellular compartments, non-aqueous fractionation of organelles was performed. No significant differences between the genotypes were observed (Fig. S1F), suggesting that the mechanisms by which RCD1 regulates chloroplastic redox status are not dependent on the photosynthetic ETC or nucleotide electron carriers.

### Mitochondrial AOX respiration and energy metabolism are altered in *rcd1*

In summary, the experiments revealed defects in *rcd1* chloroplasts which could not be explained by any of the tested chloroplast functions. Earlier transcriptomic studies in *rcd1* did not reveal marked changes in the expression of chloroplast components which could affect photosynthetic ETC. However, these studies did reveal that a number of genes encoding mitochondrial functions, including *AOX,* had increased expression in *rcd1* under standard growth conditions (Jaspers et al. 2009, Brosche et al. 2014). Immunoblotting of isolated mitochondria with an antibody recognizing all five isoforms of Arabidopsis AOX (AOX1a, -b, -c, -d, and AOX2) confirmed increased abundance of AOX in *rcd1* (Fig. 2A). The most abundant AOX isoform in Arabidopsis is AOX1a. Accordingly, only a very weak signal was detected in the *aox1a* mutant. However, in the *rcd1 aox1a* double mutant (Brosché et al. 2014) AOXs other than AOX1a were evident, thus dysfunction or absence of RCD1 led to the increased abundance of several AOX isoforms.

**Figure 2.**
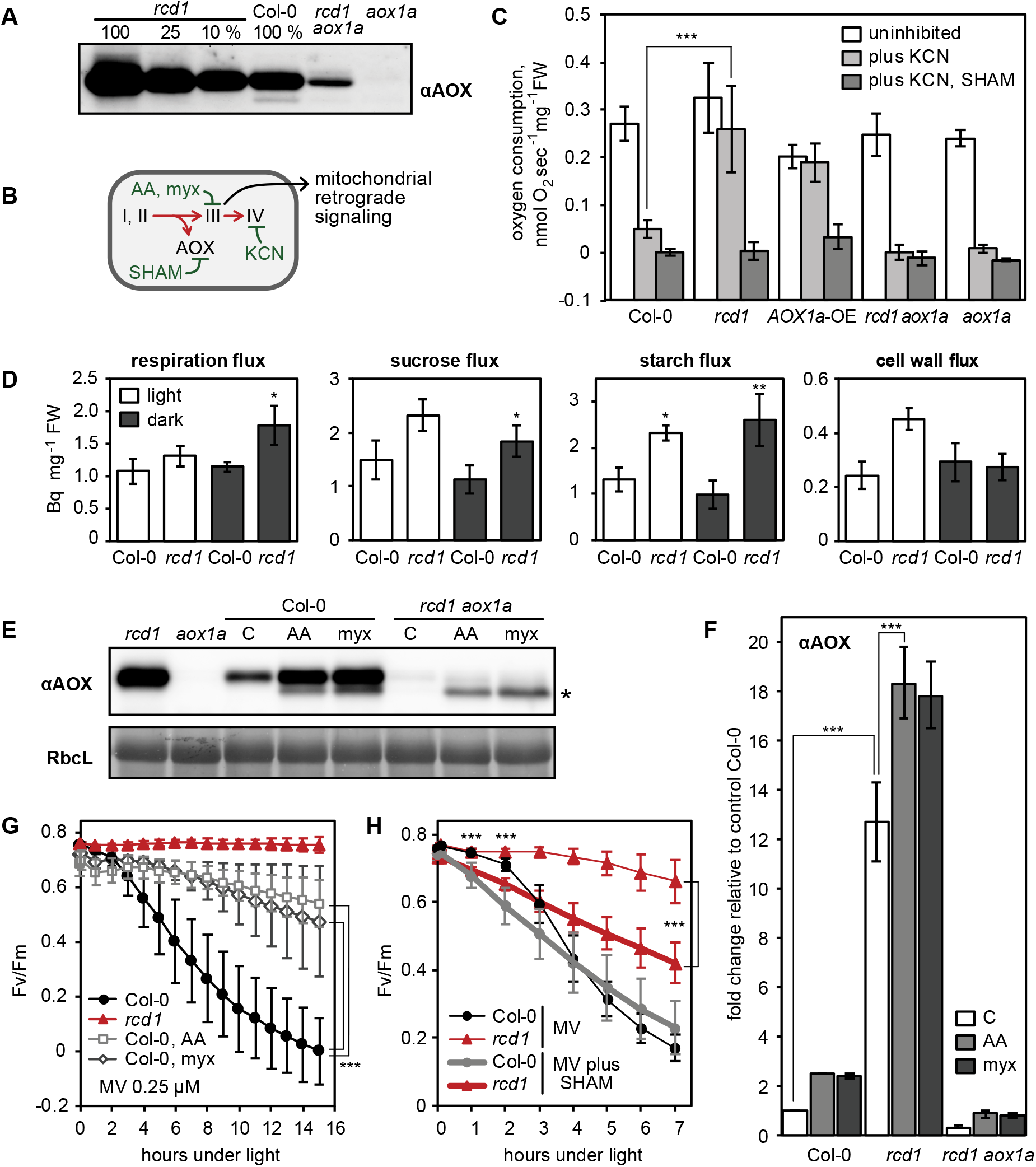
Mitochondrial AOXs affect energy metabolism of *rcd1* and alter response to chloroplastic ROS. (A) Abundance of AOX isoforms in mitochondrial preparations, as assessed with αAOX antibody. (B) Two mitochondrial respiratory pathways (red arrows) and sites of action of mitochondrial inhibitors (green). KCN inhibits complex IV. SHAM inhibits AOX activity. Antimycin A (AA) and myxothiazol (myx) block electron transfer through complex III, creating ROS-related mitochondrial retrograde signal. (C) Oxygen uptake by seedlings in the darkness in presence of mitochondrial respiration inhibitors [mean ± SD, asterisks denote selected values that are significantly different (P value < 0.001, one-way ANOVA with bonferroni post hoc correction, for full statistical analysis see Supplementary Table 5)]. (D) Deduced metabolic fluxes in light- and dark-adapted Col-0 and *rcd1* assessed by fractionation of [U-^14^C] glucose-labelled leaf extracts (mean ± SE, asterisks indicate values significantly different from the wild type, **P value < 0.01, *P value < 0.05, Student’s t-test). (E) Changes in AOX abundance after pretreatment with AA or myx (C – control treatment with no inhibitor). Putative position of AOX1d is labeled with an asterisk. (F) Quantification of αAOX immunostaining signal after pretreatment with AA or myx (C – control treatment with no inhibitor). To avoid saturation of αAOX signal in *rcd1,* a dilution series of protein extracts was made [mean ± SD, asterisks denote selected values that are significantly different (P value < 0.001, bonferroni post hoc correction, for full statistical analysis see Supplementary Table 5)]. (G) MV-induced PSII photoinhibition in leaf discs treated as in panels E and F [mean ± SD, asterisks indicate selected treatments that are significantly different (P value < 0. 001, bonferroni post hoc correction, for full statistical analysis see Supplementary Table 5)]. (H) Effect of SHAM on PSII photoinhibition by chloroplastic ROS [mean ± SD, asterisks indicate significant difference in the treatments of the same genotype at the selected time points (P value < 0.001, bonferroni post hoc correction, for full statistical analysis see Supplementary Table 5)].

In order to test whether the high abundance of AOXs in *rcd1* correlated with their increased activity, seedling respiration was assayed *in vivo.* Mitochondrial AOXs form an alternative respiratory pathway to the KCN-sensitive electron transfer through complex III and cytochrome C (Fig. 2B). Thus, after recording the initial rate of O_2_ uptake, KCN was added to inhibit cytochrome-dependent respiration. In Col-0 seedlings KCN led to approximately 80 *%* decrease in O_2_ uptake, *versus* only about 20 *%* in *rcd1* (Fig. 2C). Elevated AOX capacity of *rcd1* was similar to that of an *AOX1a-OE* overexpressor line expressing *AOX1a* under CMV 35S promoter (Umbach et al., 2005). In *rcd1 aox1a* AOX capacity was the same as in Col-0 or *aox1a* (Fig. 2C). Thus, elevated AOX respiration of *rcd1* was largely dependent on the AOX1a isoform.

The ATP yield of the AOX-dependent respiration is significantly lower than that of the cytochrome-dependent pathway (Vanlerberghe et al. 2016). Thus, continuously induced AOX activity is likely to cause changes in cell’s energy metabolism. To test whether this was the case in *rcd1,* leaf discs from light- and dark-adapted plants were treated with ^14^C-labeled glucose, and distribution of radioactive label between evolved ^14^CO_2_ and fractionated plant material was analyzed. Significantly increased carbohydrate metabolism was detected in *rcd1* (Fig. 2D, Supplementary Table 1). The redistribution of radiolabel to sucrose, starch and cell wall was elevated in *rcd1* as were the corresponding deduced fluxes (Fig. 2D). Increased respiratory flux and higher amount of total metabolized glucose (Supplementary Table 1) in *rcd1* indicated more active glycolytic pathway. Furthermore, higher cell wall metabolic flux observed in *rcd1* provided indirect support of elevated operation of the oxidative pentose phosphate pathway which is required for generating pentans used in cell wall biosynthesis (Ap Rees 1978). Unfortunately, the discontinued commercial availability of the required positionally radiolabeled glucoses prevented us from analyzing fermentative fluxes more directly. Interestingly, metabolism of the *rcd1 aox1a* mutant was only slightly different from *rcd1* under light and indistinguishable from *rcd1* in the darkness (Supplementary Table 1). Taken together, these results suggested that inactivation of *RCD1* led to marked changes in energy metabolism likely including activation of glycolytic and oxidative pentose phosphate pathways. Similar alterations observed in the *rcd1* and the *rcd1 aox1a* mutants indicated that the AOX1a isoform was not playing a decisive role in these changes.

### Mitochondrial AOXs contribute to ROS processing in the chloroplasts

The inhibition of complex III by AA or myx activates mitochondrial retrograde signaling (Fig. 2B), leading to nuclear transcriptional reprogramming including induction of *AOX* genes (Clifton et al., 2006). Accordingly, treatment with either of these chemicals significantly increased abundance of AOXs in all tested lines (Fig. 2E, F). The *rcd1 aox1a* double mutant accumulated AOXs other than AOX1a, including a putative AOX1d (Konert et al., 2015) (Fig. 2E). Treatment of *rcd1* with AA or myx led to a further increase in AOX abundance despite high steady state levels (Fig. 2F). This showed that responsiveness of *rcd1* to the complex III retrograde signal was not compromised, but rather appeared to be continuously active.

To assess whether increased AOX abundance correlated with changes in chloroplast functions, PSII inhibition was assayed in the presence of MV in AA-or myx-pretreated leaf discs. Pretreatment of Col-0 with any of the chemicals increased resistance of PSII to inhibition by chloroplastic ROS (Fig. 2G). In addition to complex III, AA has been reported to inhibit plastid cyclic electron flow by binding to PGR5 (PROTON GRADIENT REGULATION 5) (Sugimoto et al., 2013). This possible off-target effect was assessed by using the *pgr5* mutant. Pretreatment with AA made both *pgr5* and its background wild type *gl1* equally more tolerant to chloroplastic ROS (Fig. S2A), showing that an effect on PGR5 was not the reason for the observed gain in ROS tolerance.

Mitochondrial retrograde signaling triggered by AA and myx (Fig. 2B) induces expression of several genes other than AOX. To test whether accumulation of AOXs contributed to PSII protection from chloroplastic ROS or merely correlated with it, the AOX inhibitor salicylhydroxamic acid (SHAM) was used. Treatment of plants with SHAM alone resulted in very mild PSII inhibition, which was identical in *rcd1* and Col-0 (Fig. S2B). However, pretreatment with SHAM made both *rcd1* and Col-0 plants significantly more sensitive to chloroplastic ROS generated by MV (Fig. 2H). Another potential target of SHAM is PTOX, the terminal oxidase of the chloroplast that is analogous to AOX (Fu et al., 2012). To assess involvement of PTOX, the response of the *ptox* mutant to the same treatment as above was tested. As in Col-0, in the *ptox* mutant MV plus SHAM resulted in a more rapid decline in PSII activity than MV alone (Fig. S2C), suggesting that PTOX was not involved in the observed decrease of ROS tolerance. Taken together, these results indicated that mitochondrial AOXs contributed to resistance of PSII to chloroplastic ROS.

To address the role of the AOX1a isoform in this process, MV-induced PSII inhibition was tested in the *AOX1a-OE* line (Umbach et al. 2005) and in the *aox1a* mutant. Immunoblotting showed that abundance of AOX protein in *AOX1a-OE* was comparable to that in *rcd1* (Fig. S2D). However, tolerance of either *AOX1a-OE* or *aox1a* plants to chloroplastic ROS was indistinguishable from the wild type (Fig. S2E). Thus, AOX1a overexpression was not sufficient to provide ROS tolerance. When the analogous experiment was performed in *rcd1 aox1a* – despite exhibiting lower levels of AOX proteins (Fig. 2A, E, F) and a wild-type AOX respiration capacity (Fig. 2C), *rcd1 aox1a* was as tolerant to MV as *rcd1* (Fig. S2F). These results suggest that AOX isoforms other than AOX1a are likely to contribute to the tolerance of *rcd1* to chloroplastic ROS. Moreover, AOX activity appeared to be only part of the molecular mechanism required to affect chloroplast ROS processing.

### Evidence for increased electron transfer between chloroplasts and mitochondria in *rcd1*

The above-described results indicated that mitochondrial AOXs affected ROS processing in the chloroplasts. As this process spans both mitochondria and chloroplasts, it seems reasonable to suggest that some of its components are related to electron transfer between the two organelles. Chlorophyll fluorescence under light (Fs) was used as a proxy of the reduction state of plastoquinone, one of the electron carriers of the chloroplast ETC. After combined treatment with SHAM and MV (as in Fig. 2H), Fs increased in *rcd1,* but not in Col-0 (Fig. 3A). This suggests that a pathway is activated in *rcd1* linking the photosynthetic ETC to the activity of mitochondrial AOXs, with the latter functioning as electron sink. When the AOX activity was inhibited by SHAM, electron flow along this pathway was blocked, leading to an accumulation of electrons in the chloroplast ETC.

**Figure 3.**
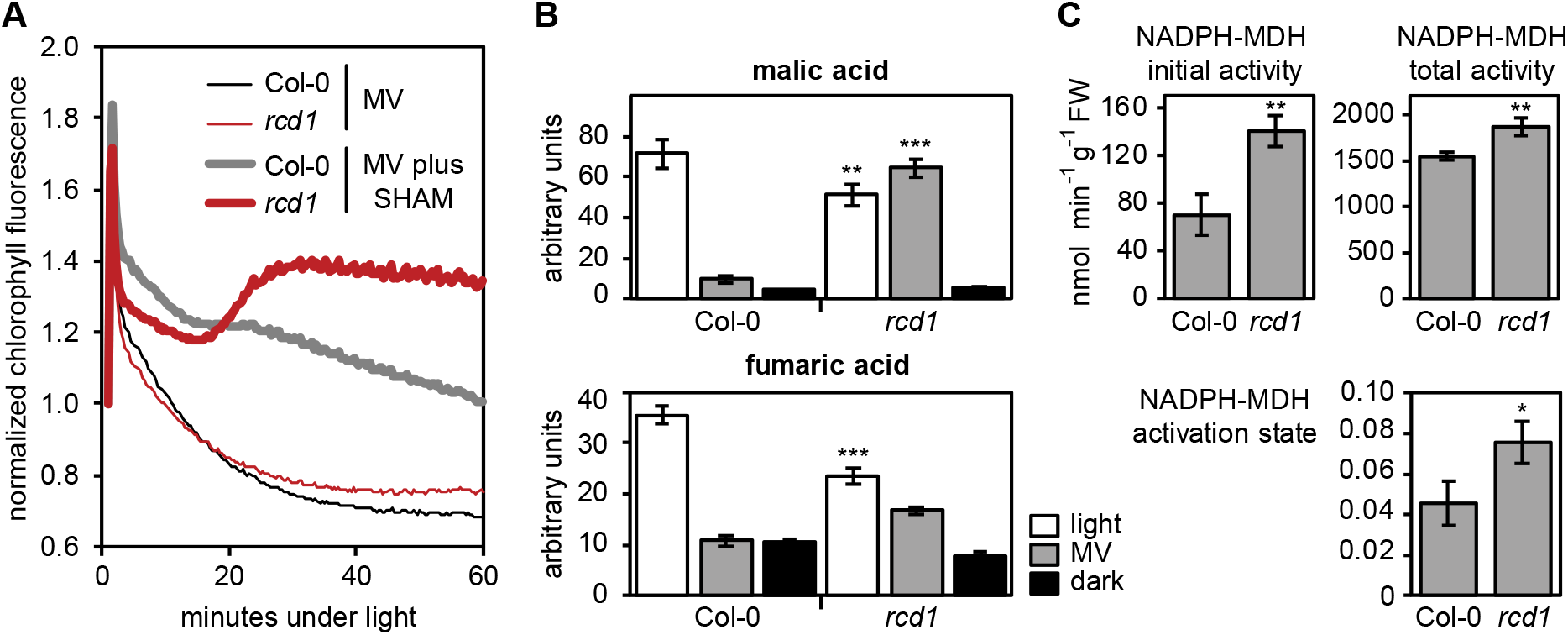
Altered electron transfer between the organelles in *rcd1.* (A) Effect of MV and SHAM treatment on chlorophyll fluorescence under light. (B) Levels of malic and fumaric acids in extracts from Col-0 and *rcd1* seedlings that were either exposed to light in absence or in presence of MV, or were dark-adapted (mean ± SE, asterisks indicate values significantly different from those in the similarly treated wild type, ***P value < 0.001, **P value < 0.01, Student’s t-test). (C) Activity of chloroplastic NADPH-MDH malate dehydrogenase assessed without thiol-reducing agent DTT (initial activity, top left) and in presence of DTT (total activity, top right). The activation state of NADPH-MDH (bottom) is presented as the ratio of the initial and the total activity (mean ± SE, asterisks indicate values significantly different the wild type, **P value < 0.01, *P value < 0.05, Student’s t-test).

One of the mediators of electron transfer between the organelles is malate (Scheibe 2004, Zhao et al. 2018). Under light, some of the reduced NADPH generated by the photosynthetic ETC is used by the chloroplast NADPH-dependent malate dehydrogenase (NADPH-MDH) that reduces oxaloacetate to malate. Malate is subsequently exported from chloroplasts, concomitantly mediating the export of reducing equivalents. Levels of malate were assessed in total extracts from Col-0 and *rcd1* seedlings which were either dark-adapted overnight, or exposed to 4 hours of light in the presence or absence of MV. Treatment of Col-0 with MV induced a significant decrease in malate levels; however, this change was not apparent following treatment of *rcd1* with MV (Fig. 3B). It seems likely that this effect is specific to malate, given that alterations in the level of the related metabolite fumarate did not show opposing response to MV treatment in the two genotypes (Fig. 3B).

The activity of NADPH-MDH was next tested in extracts from the light-adapted plants. Chloroplast NADPH-MDH is a redox-regulated enzyme activated by reduction of thiol bridges. Thus, the NADPH-MDH activity may reflect the *in vivo* thiol redox state of the cellular compartment from which it has been isolated. Hence, activity was first assessed without added thiol reductants (initial activity, Fig. 3C), and then DTT was added to fully activate the enzyme (total activity, Fig. 3C). Both values were higher in *rcd1* than in Col-0. To determine the contribution of *in vivo* thiol redox state, the initial NADPH-MDH activity was divided by total activity. This value, the activation state, also increased in *rcd1* (Fig. 3C). An increased activation state of NADPH-MDH was consistent with a more reduced thiol redox status of *rcd1* chloroplasts as observed in the analyses of 2-CP oxidation (Fig. 1E). In addition, the increased total activity of NADPH-MDH in *rcd1* indicated that regulatory factors other than the thiol status also contributed to its higher activity in *rcd1.* Taken together, these results suggest that mitochondria contributed to ROS processing in the chloroplasts *via* a mechanism involving mitochondrial AOXs and malate shuttling and that RCD1 is involved in the regulation of these processes.

### Retrograde signaling from both chloroplasts and mitochondria is altered in *rcd1*

Our results demonstrated that absence or dysfunction of RCD1 causes physiological alterations in both chloroplasts and mitochondria. As RCD1 is a nuclear-localized transcriptional co-regulator (Jaspers et al. 2009, Jaspers et al. 2010), its involvement in retrograde signaling pathways from both organelles was assessed. For this purpose, transcriptional changes observed in the *rcd1* mutant (Jaspers et al. 2009, Brosche et al. 2014) were compared to other published transcriptomic changes caused by perturbations in energy organelles. These analyses revealed a striking similarity of genes differentially regulated in *rcd1* to those affected by disturbed organellar function (Fig. S3). Among the analyzed perturbations were disruptions of mitochondrial genome stability *(msh1 recA3),* organelle translation *(mterf6, prors1),* activity of mitochondrial complex I *(ndufs4,* rotenone), complex III (AA), and ATP synthase function (oligomycin), as well as treatments and mutants related to chloroplastic ROS production (high light, MV, H_2_O_2_, *alx8/ fry1*, norflurazon).

In particular, a significant overlap was observed between genes mis-regulated in *rcd1* and the mitochondrial dysfunction stimulon (MDS) genes (De Clercq et al. 2013) as well as with genes affected by 3’-phosphoadenosine 5’-phosphate (PAP) signaling (Estavillo et al. 2011, Van Aken and Pogson 2017) (Fig. 4A). Given that PAP signaling is suppressed by the activity of SAL1, it was increased in the mutants deficient in SAL1 *(alx8* and *fry1,* Fig. 4A, S3). As expected, *AOX1a* was among the genes induced by the majority of the treatments. Another MDS gene with increased expression in *rcd1* encoded the sulfotransferase SOT12, an enzyme generating PAP. Accordingly, immunoblotting of total protein extracts with αSOT12 antibody demonstrated elevated SOT12 protein abundance in *rcd1* (Fig. 4B). To address the functional interaction of RCD1 with PAP signaling, *rcd1-4* was crossed with *sal1-8* (also known as *alx8).* The resulting *rcd1 sal1* mutant was severely affected in development (Fig. 4C). Together with transcriptomic similarities between *rcd1* and *sal1* mutants, this further supported on the overlap of PAP and RCD1 signaling pathways.

**Figure 4.**
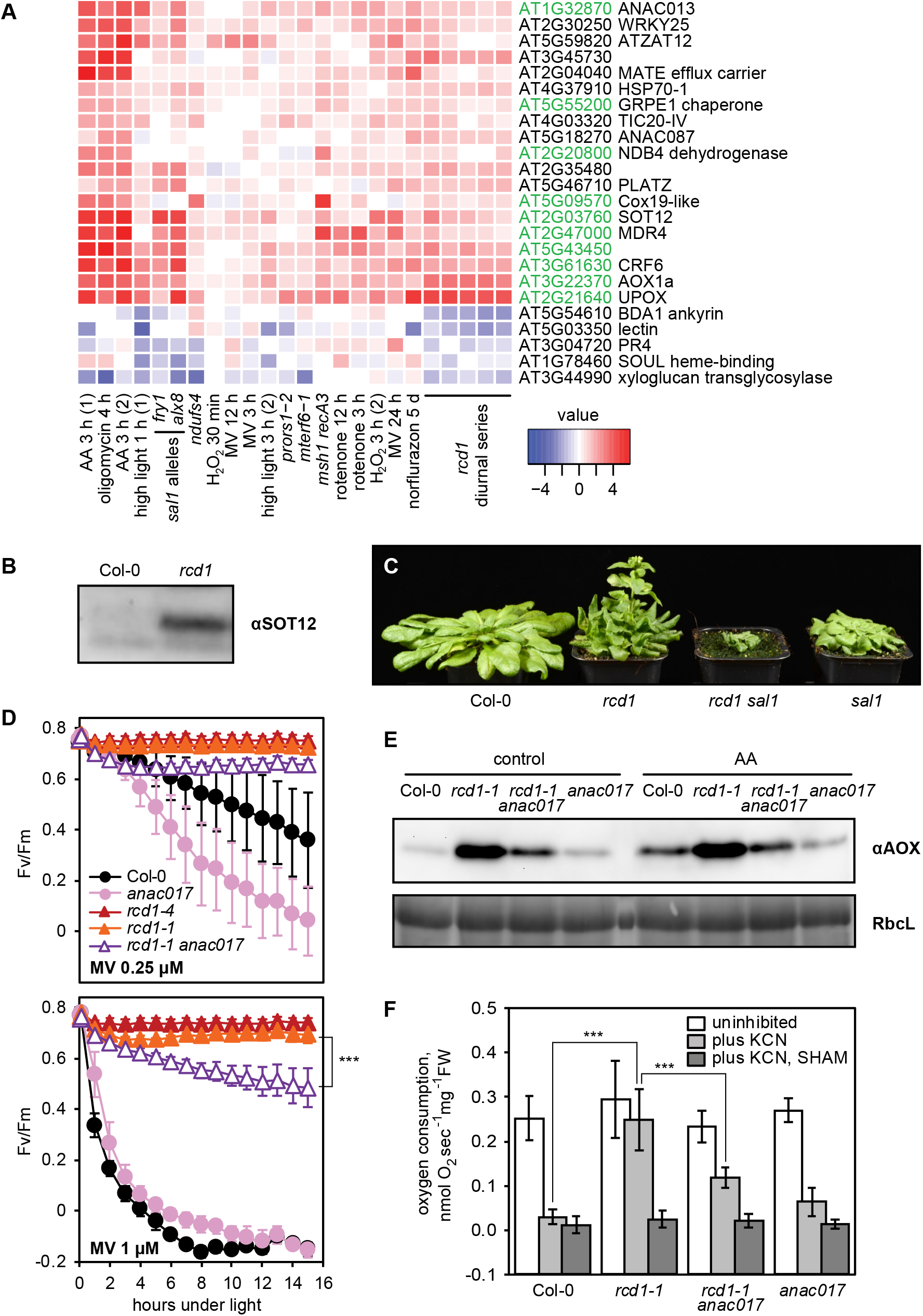
RCD1 is involved in chloroplast ROS and mitochondrial ANAC017 retrograde signaling pathways. (A) Regulation of *rcd1* mis-expressed genes under perturbations of organellar functions in the selected subset of genes. The full unbiased list of rcd1-misexpressed genes is presented in Fig. S3. MDS genes are labeled green. (B) Sulfotransferase SOT12 encoded by an MDS gene accumulates in *rcd1* under standard growth conditions, as revealed by immunoblotting with the specific antibody. (C) Phenotype of *rcd1 sal1* double mutant under standard growth conditions. (D) PSII photoinhibition by ROS in *rcd1 anac017* double mutant, triggered with 0.25 μM or 1 μM MV (top and bottom panel, accordingly) [mean ± SD, asterisks denote values significantly different from those in the similarly treated wild type at the last time point of the assay (P value < 0.001, two-way ANOVA with bonferroni post hoc correction, for full statistical analysis see Supplementary Table 5)]. (E) Total AOX protein levels in *rcd1 anac017* double mutant. (F) Oxygen uptake by *rcd1 anac017* seedlings in the darkness in presence of mitochondrial respiration inhibitors [mean ± SD, asterisks denote selected values that are significantly different (P value < 0.001, one-way ANOVA with bonferroni post hoc correction, for full statistical analysis see Supplementary Table 5)].

Expression of MDS genes is regulated by the transcription factors ANAC013 and ANAC017 (De Clercq et al. 2013). To elucidate whether genes targeted by ANAC013/ ANAC017 were enriched among genes with altered mRNA abundance in the *rcd1* mutant, the presence of the ANAC-responsive cis-element CTTGNNNNNCA[AC]G (De Clercq et al. 2013) was assessed in promoter regions of *rcd1* mis-regulated genes. Significant enrichment of this cis-element was found (Fig. S3). Increased expression of several MDS genes in *rcd1* together with the fact that RCD1 interacted with ANAC013 *in vitro* and in the yeast two-hybrid system (Jaspers et al. 2009, O’Shea et al., 2015, O’Shea et al., 2017) suggests that RCD1 is a negative regulator of ANAC013/ ANAC017 transcription factors. It should be remembered that MDS genes represented only a subclass of all genes whose expression is affected or co-regulated by RCD1 (Fig. S3). For example, a cluster of genes that have lower expression in both *rcd1* and *sal1* mutants and are mostly associated with defense against pathogens did not have enrichment of ANAC motif in their promoters (Fig. 4A, S3). This is likely a consequence of interaction of RCD1 with about forty different transcription factors belonging to several families (Jaspers et al. 2009).

Physiological outcomes of the interaction between RCD1 and ANAC transcription factors were tested by reverse genetics. ANAC017 has been shown to regulate the expression of *ANAC013* in the mitochondrial retrograde signaling cascade (Van Aken et al. 2016). Since no *anac013* knockout mutant is available, only the *rcd1-1 anac017* double mutant was generated. It was more sensitive to chloroplastic ROS than the parental *rcd1* line (Fig. 4D). In addition, *rcd1 anac017* was characterized by lower abundance of AOX isoforms (Fig. 4E) and reduced AOX respiration capacity (Fig. 4F) compared to *rcd1.* Thus, both chloroplast- and mitochondria-related phenotypes of *rcd1* were partially mediated by ANAC017.

### RCD1 interacts with ANAC transcription factors *in vivo*

To assess whether RCD1 associates with ANAC transcription factors *in vivo,* two independent pull-down experiments were carried out. To identify interaction partners of ANAC013, an Arabidopsis line expressing ANAC013-GFP (De Clercq et al. 2013) was used. ANAC013-GFP was purified with αGFP beads, and associated proteins were identified by mass spectrometry in three replicates. RCD1, its closest homolog SRO1, as well as ANAC017 were identified as ANAC013 interacting proteins; see Table 1 for a list of selected nuclear-localized interaction partners of ANAC013, and Supplementary Table 2 for the full list of identified proteins and mapped peptides. These data confirmed that ANAC013, RCD1 and ANAC017 are components of the same protein complex *in vivo.* In a reciprocal pull-down assay using transgenic line expressing RCD1 tagged with triple Venus YFP under the control of *UBIQUITIN10* promoter, RCD1-3xVenus and interacting proteins were immunoprecipitated using αGFP (Table 1; Supplementary Table 3). Transcription factor ANAC017 was found among RCD1 interactors.

**Table 1.** Overview of the immunoprecipitation results. Selected proteins identified in ANAC013-GFP and RCD1-3xVenus pull-down assays. Ratio *vs.* Col-0 and the P-value were obtained by Perseus statistical analysis from the three repeats for each genotype used. Bold text indicates baits. The peptide coverage for selected proteins as well as full lists of identified proteins are presented in Supplementary Tables 2 and 3.

| ***ANAC013-GFP pull-down*** |  |  |  |  |  |
| --- | --- | --- | --- | --- | --- |
| Ratio ANAC013 vs. Col-0 | P-value | unique peptides | gene | name | stickiness |
| **50966** | **7.09 x 10<sup>-7</sup>** | **29** | **AT1G32870** | **ANAC013** |  |
| **22149** | **3.41 x 10<sup>-8</sup>** | **25** |  | **GFP** |  |
| 10097 | 3.67 x 10<sup>-6</sup> | 37 | AT1G32230 | RCD1 | 1.00 % |
| 110 | 1.67 x 10<sup>-6</sup> | 8 | AT2G35510 | SRO1 | 1.00 % |
| 74 | 1.09 x 10<sup>-9</sup> | 4 | AT1G34190 | ANAC017 | 1.00 % |
| ***RCD1-3xVenus pull-down*** |  |  |  |  |  |
| Ratio RCD1 vs. Col-0 | p-value | unique peptides | gene | name | stickiness |
| **7593** | **0.000454** | **35** | **AT1G32230** | **RCD1** |  |
| **1292** | **0.006746** | **10** |  | **YFP** |  |
| 108 | 5.48 x 10<sup>-8</sup> | 2 | AT1G34190 | ANAC017 | 1.00 % |

To test whether RCD1 directly interacts with ANAC013/ ANAC017 *in vivo,* the complexes were reconstituted in the human embryonic kidney cell (HEK293T) heterologous expression system. RCD1 tagged with N-terminal triple HA (HA-RCD1) and ANAC013 tagged with C-terminal triple myc (ANAC013-myc) were co-expressed in HEK293T cells. Co-immunoprecipitation of ANAC013-myc with αHA and of HA-RCD1 with αmyc (Fig. 5A) confirmed complex formation between the two proteins. Then a similar experiment was performed for the interaction pair RCD1-ANAC017. Co-immunoprecipitation of ANAC017-myc with αHA and of HA-RCD1 with αmyc confirmed complex formation between the two proteins co-expressed in HEK293T cells (Fig. 5B). Together with the results of *in vivo* pull-down assays, these experiments strongly indicated formation of the complex between RCD1 and ANAC013/ ANAC017 transcription factors.

**Figure 5.**
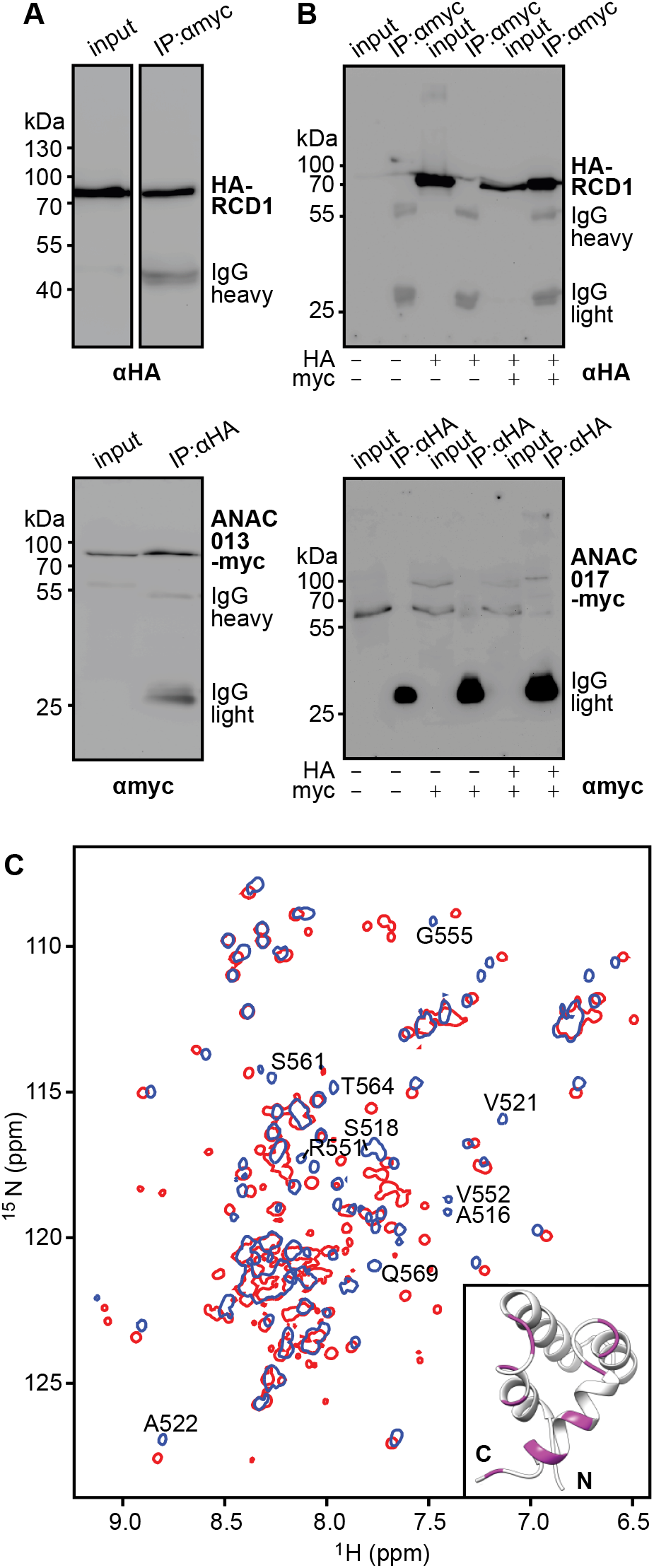
Biochemical interaction of ANAC013/ ANAC017 transcription factors with the RST domain of RCD1. HA-RCD1 was co-expressed with ANAC013-myc (A) or ANAC017-myc (B) in human embryonic kidney cells (IP – eluate after immunoprecipitation). Interaction of the C-terminal domain of RCD1 with ANAC013-derived peptide caused peptide-induced chemical shift changes in the ^1^H, ^15^N correlation spectrum of RCD1^468-589^, which were mapped on the structure of the RST domain (C). (A) Co-immunoprecipitation of HA-RCD1 with αmyc antibody (top) and of ANAC013-myc with αHA-antibody (bottom). (B) Co-immunoprecipitation of HA-RCD1 with αmyc antibody (top) and of ANAC017-myc with αHA-antibody (bottom). (C) Superimposed ^1^H, ^15^N HSQC spectra of the C-terminal domain of RCD1 acquired in absence (blue) and presence (red) of approximately two-fold excess of the ANAC013^235-284^ peptide. Inset: RS_TRCD1_ structure with highlighted residues demonstrating the largest chemical shift perturbations between the free and bound forms (details in Fig. S4D).

### Structural insights into RCD1-ANAC interactions

RCD1 interacts with many transcription factors belonging to different families (Jaspers et al. 2009, Jaspers et al. 2010). The interaction is mediated by the RCD1 C-terminal RST domain. The strikingly diverse set of RCD1 interacting partners may be partially explained by disordered flexible regions present in the transcription factors (Kragelund et al., 2012, O’Shea et al. 2017). The C-terminal domain of RCD1 (residues 468-589) including the RST domain (RST_RCD1_; 510-568) was purified and labeled with ^13^C and ^15^N for NMR spectroscopic study (Tossavainen et al., 2017). The solution NMR structure of the C-terminal domain and the structural statistics for the ensemble of the 15 lowest-energy structures of RCD1^468-589^ including the RST domain are presented in Fig. S4A and Supplementary Table 4, respectively. The atomic coordinates and structural restraints for the C-terminal domain of RCD1^468-589^ have been deposited in the Protein Data Bank with the accession code 5N9Q. The RS_TRCD1_ is entirely α-helical, with unstructured flanking regions. The structured region ends with position N568, which corresponds to the necessary C-terminal part for the interaction with transcription factors (Jaspers et al. 2010).

ANAC013 has been shown to interact with RCD1 in yeast-two-hybrid assays (Jaspers et al. 2009, O’Shea et al. 2017), in which interaction with RCD1 was mapped to ANAC013 residues 205-299 (O’Shea et al. 2017). Thus, we selected ANAC013 to address the specificity of the interaction of the RST domain upon interaction with transcription factors. Screening of ANAC013 peptides by surface plasmon resonance (details in Fig. S4B, C) revealed the ANAC013^235-284^ peptide as the strongest RCD1 interactor, thus this peptide was chosen for further NMR analysis with purified C-terminal domain of RCD1. Superimposed ^1^H, ^15^N HSQC spectra of RCD1^468-589^ acquired in absence (blue) and presence (red) of approximately two-fold excess of the ANAC013^235-284^ peptide are presented in Figure 5C. Combined ^1^H, ^15^N chemical shift perturbations (CSPs) in backbone amide chemical shifts upon addition of the peptide are depicted as a histogram and mapped onto the structure of the RS_TRCD1_ domain (Fig. S4D and Fig. 5C inset, accordingly). The largest CSPs (Δδ ≥ 0.10 ppm) were found for residues located on one face of the domain, formed by the first and last helices and loops between the first and the second, and the third and the last helices. This region probably represents the peptide interaction site. However, relatively large perturbations were observed throughout the domain, and notably, in the unstructured C-terminal tail, which might originate from a conformational rearrangement in the domain induced by ligand binding. These data confirmed strong and specific binary interaction between the RCD1 RST domain and the ANAC013 transcription factor.

### RCD1 protein is sensitive to organellar ROS

RCD1 is a nuclear-localized transcriptional co-regulator that affects chloroplastic and mitochondrial functions by interacting with transcription factors of mitochondrial retrograde signaling (Figures 1–5). Thus, it was logical to assume that the RCD1 protein may be post-translationally modified in response to feedback regulation by organellar signaling. To test this, it was first assessed whether RCD1 was directly affected by chloroplast ROS *in vivo.* For this, a complemented *rcd1-4* line expressing RCD1 tagged with triple HA epitope at the C-terminus under the *RCD1* native promoter was used (Jaspers et al. 2009). No changes were observed in RCD1-HA levels during 5 hours of standard growth light amid the light period, or during 5-hour high light treatment (1 300 μmol m^−2^ s^−1^). On the other hand, both exogenous MV (1 μM) and H_2_O_2_ (100 mM) treatments led to decreased RCD1 abundance (Fig. 6A). When plant extracts from these experiments were separated in non-reducing SDS-PAGE, the RCD1-HA signal resolved into several species of different molecular weight, representing oligomeric forms of RCD1 (Fig. 6B, left panels). Under standard growth conditions and under high light treatment, most RCD1-HA formed a reduced monomer. However, treatment with MV or H_2_O_2_ resulted in fast conversion of RCD1-HA into high-molecular-weight aggregates (Fig. 6B, left panels).

**Figure 6.**
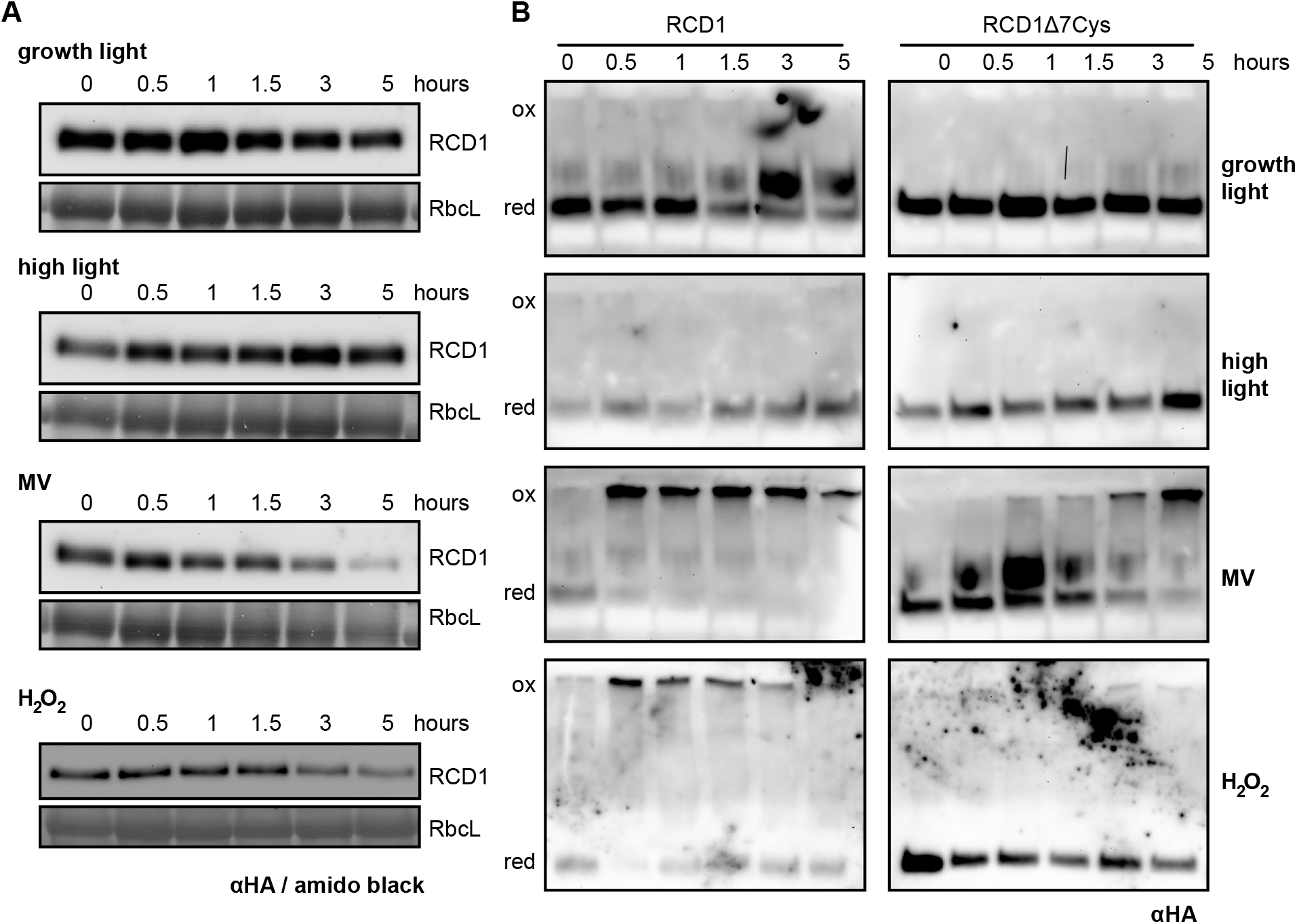
RCD1 protein is sensitive to chloroplastic ROS. (A) Plants expressing HA-tagged RCD1 under native promoter were treated with 5-hour growth light (150 μmol m^−2^ s^−1^), high light (1 300 μmol m^−2^ s^−1^), MV (1 μM) or H_2_O_2_ (100 mM). The level of RCD1-HA protein was monitored by immunoblotting with αHA antibody at different time points as indicated. Rubisco large subunit (RbcL) amounts detected by amido black staining are shown in all panels as a control for equal protein loading. (B) Separation of total protein extracts in non-reducing PAGE to reveal thiol-bonded forms of RCD1 in wild-type RCD1-HA and in the RCD1Δ7Cys mutant. Reduced (red) and oxidized (ox) forms of the protein are labeled.

To investigate whether RCD1 aggregates were thiol-dependent, a complementation line was created that expressed an RCD1 variant in which the cysteines in the linkers between the RCD1 domains (7 cysteine residues out of 13 in total) were mutated to alanines (RCD1Δ7Cys = RCD1 C14A-C37A-C50A-C175A-C179A-C212A-C243A; Fig. S5A). The chemical treatments performed in the RCD1Δ7Cys-HA line caused significantly less aggregation of RCD1 (Fig. 6B, right panels), as well as an overall increased abundance of the monomer RCD1 form (Fig. 6B, S5B). Noteworthy, no effect on RCD1 redox state was observed under AA treatment (Fig. S5B, top panel), suggesting that the RCD1 protein was not a direct target of the mitochondrial ROS signal. Analysis of RCD1Δ7Cys levels in reducing gels demonstrated that this variant was protected from degradation by H_2_O_2_ or MV (Fig. S5B). However, the RCD1Δ7Cys-HA construct complemented the *rcd1* MV phenotype similarly to RCD1-HA (Fig. S5C). These results demonstrated that RCD1 thiols were sensitive to ROS generated in chloroplasts. Chloroplastic ROS formation led to aggregation of the nuclear RCD1 in complexes most probably stabilized by disulfide bonds. However, inhibition of RCD1 aggregation by mutating its cysteines did not change tolerance of plants to fast PSII photoinhibition triggered by MV.

## Discussion

Plant chloroplasts and mitochondria work together to supply the cell with energy and metabolites, creating a complex communication network. ROS are formed in these organelles as by-products of their electron transfer chains (photosynthetic in chloroplasts and respiratory in mitochondria). ROS serve as versatile signaling molecules regulating many aspects of plant physiology such as development, stress signaling, systemic responses, and programmed cell death (PCD) (Dietz et al. 2016, Noctor et al. 2017, Waszczak et al. 2018). This communication network also affects gene expression in the nucleus where numerous signals are perceived and integrated. The mitochondrial and chloroplastic retrograde signaling pathways have been extensively studied independently; however, molecular mechanisms of coordinated action of the two types of energy organelles in response to environment are poorly understood. Substantial evidence accumulated in this and earlier studies revealed the nuclear protein RCD1 as a central player in communication of the energy organelles with the nuclear gene expression apparatus.

The *rcd1* mutant has been characterized by alterations in both chloroplasts and mitochondria (Fujibe et al. 2004, Heiber et al. 2007, Jaspers et al. 2009, Brosché et al. 2014, Hiltscher et al. 2014), and transcriptomic outcomes of RCD1 inactivation share similarities with those triggered by disrupted functions of both organelles (Fig. 4, S3). The results here suggest that RCD1 forms inhibitory complexes with components of mitochondrial retrograde signaling *in vivo* and that redox state and stability of RCD1 are directly influenced by chloroplastic ROS. This places RCD1 into a regulatory system encompassing mitochondrial complex III signaling through ANAC013/ ANAC017 transcription factors and chloroplastic signaling by H_2_O_2_/ PAP. The existence of such inter-organellar regulatory system has been previously proposed on the basis of the results of transcriptomic analyses (Van Aken and Pogson 2017), but the molecular mechanisms underlying it were unknown. Our results place RCD1 at the intersection of these mitochondrial and chloroplast signaling pathways.

RCD1 was earlier proposed to be a transcriptional co-regulator because of its interaction with many transcription factors in yeast-two-hybrid analyses (Jaspers et al. 2009). The direct *in vivo* interaction of RCD1 with ANAC013 and ANAC017 revealed in this study (Table 1, Fig. 5) allows RCD1 to affect expression of the MDS: a set of nuclear genes activated by ANAC013/ ANAC017 and mostly encoding mitochondrial components (De Clercq et al. 2013). *ANAC013* itself is an MDS gene, thus mitochondrial signaling through ANAC013/ ANAC017 creates a self-amplifying loop. Physiological and transcriptomic data supports the role of RCD1 as a negative regulator of these transcription factors (Fig. 4). Thus, RCD1 is likely involved in the down-regulation of the ANAC013/ ANAC017 self-amplifying loop and in preventing excessive expression of MDS genes under unstressed conditions.

A negative co-regulator must itself be tightly regulated and quickly inactivated under stress conditions to allow the transcriptional system to induce target genes in response to stress. Accordingly, the RCD1 protein was sensitive to treatments triggering or mimicking chloroplastic ROS production (such as MV and H_2_O_2_) both in the short and long term. MV and H_2_O_2_ treatment of plants resulted in rapid oligomerization of RCD1 as confirmed with Western blot analyses (Fig. 6). Moreover, the level of RCD1 gradually decreased during prolonged (5 hours) stress treatments. This suggests several independent modes of RCD1 regulation at the protein level. This complicated post-translational regulation renders RCD1 analogous to another prominent transcriptional coregulator protein NPR1, a master immune regulator in Arabidopsis. NPR1 exists as a high molecular weight oligomer stabilized by intermolecular disulfide bonds between conserved cysteine residues. Accumulation of salicylic acid and cellular redox changes lead to the reduction of cysteines and release of NPR1 monomers that move into the nucleus to activate expression of defense genes (Kinkema et al., 2000, Mou et al., 2003, Withers and Dong 2016).

Similar to NPR1, the RCD1 protein has a bipartite nuclear localization signal. In addition, it is predicted to possess a nuclear export signal between the WWE and PARP-like domains. Like NPR1, RCD1 has several conserved cysteine residues. This could allow redox-controlled translocation of RCD1 between the nucleus and the cytoplasm, thus providing an additional mode of regulation, which will be addressed in further studies. Changes of RCD1 localization between the nucleus and the cytosol in response to stress conditions have previously been described in transient expression systems (Katiyar-Agarwal et al., 2006). Interestingly, mutation of seven interdomain cysteines in RCD1 largely eliminated the fast *in vivo* effect of chloroplastic ROS on redox state and stability of RCD1; however, it did not significantly alter the plant response to MV (Fig. 6, S5). This suggests that redox-dependent oligomerization of RCD1 serves to fine-tune its activity, while the resistance of the *rcd1* mutant to MV may be the result of a long-term developmental adaptation associated with continuous expression of MDS genes in *rcd1.*

How induction of MDS genes contributes to the energetic and signaling landscape of the plant cell remains to be investigated. Our results suggest that one notable component of this adaptation is the activity of mitochondrial alternative oxidases (AOXs). *AOX* genes are part of the MDS regulon, thus AOX protein accumulates at higher amounts in the *rcd1* mutant and in the wild type pretreated with AA or myx (Fig. 2). In both cases, elevated AOX abundance coincided with increased tolerance to chloroplastic ROS. Moreover, treatment of either *rcd1* or wild type with the AOX inhibitor SHAM made the plants more sensitive to MV, indicating direct involvement of AOX activity in the chloroplastic ROS processing.

AOX accumulation affects different aspects of energy metabolism. Enzymatic activity of AOXs results in reduction of molecular oxygen to water by utilizing the reducing power of the mitochondrial ubiquinol pool. Thus, the direct outcome of AOX activity might be the oxidation of the ubiquinol pool (Vanlerberghe et al. 2016). In agreement with this hypothesis, *rcd1* phenotypically resembles mitochondrial mutants with oxidized ubiquinol pools that are often characterized by curly leaves (Hsieh et al., 2015)(Fig. 4C). Oxidation of ubiquinol is likely to decrease electron pressure on complex III, reducing the ROS formation by complex III and thus the mitochondrial retrograde signal that triggers expression of the MDS stimulon. This negative feedback regulation of AOX expression is disrupted in the *rcd1* mutant where ANAC013/ ANAC017 transcription factors are probably engaged in perpetual mutual activation.

The effect of increased AOX activity spreads beyond mitochondria. AOX-dependent respiration generates less ATP than the cytochrome-dependent pathway (Vanlerberghe et al. 2016). Thus, constantly induced AOX activity of the *rcd1* mutant is likely to affect its energy metabolism. In agreement with this assumption, the results indicate changes in energy metabolism of *rcd1* and increased metabolic flux through the glycolytic and the oxidative pentose phosphate pathways (Fig. 2D, Supplementary Table 1). These pathways generate reducing power, which could contribute to redox abnormalities of the *rcd1* mutant. Thus, more reduced states of certain chloroplast enzymes (Fig 1E, 3C) and changed malate levels (Fig. 3B) observed in *rcd1* may by the consequences of compensatory metabolic changes triggered by increased AOX function. In agreement with this, inhibition of AOX activity in *rcd1* coincided with the symptoms indicative of reduction of plastoquinone pool in the chloroplast ETC (Fig. 3A).

It appears that AOXs in the mitochondria form an electron sink that indirectly contributes to the oxidization of the electron acceptor side of PSI. This inter-organellar electron transfer may decrease production of ROS by PSI (Fig. 7). Altered levels of malic acid and increased activity of NADPH-dependent malate dehydrogenase in *rcd1* (Fig. 3) suggest the involvement of the malate shuttle. Accordingly, the malate shuttle was recently shown to mediate a chloroplast-to-mitochondria electron transfer pathway that caused ROS production by complex III and mitochondrial retrograde signaling (Wu et al., 2015, Zhao et al. 2018). Moreover, another inter-organellar electron transfer pathway, photorespiration, was also proposed to link chloroplastic ROS production with AOXs (Watanabe et al. 2016). It is thus possible that the chloroplast-to-mitochondrial electron transfer takes place through more than one pathway, and that the relative contribution of each of the alternatives depends on the physiological context.

**Figure 7.**
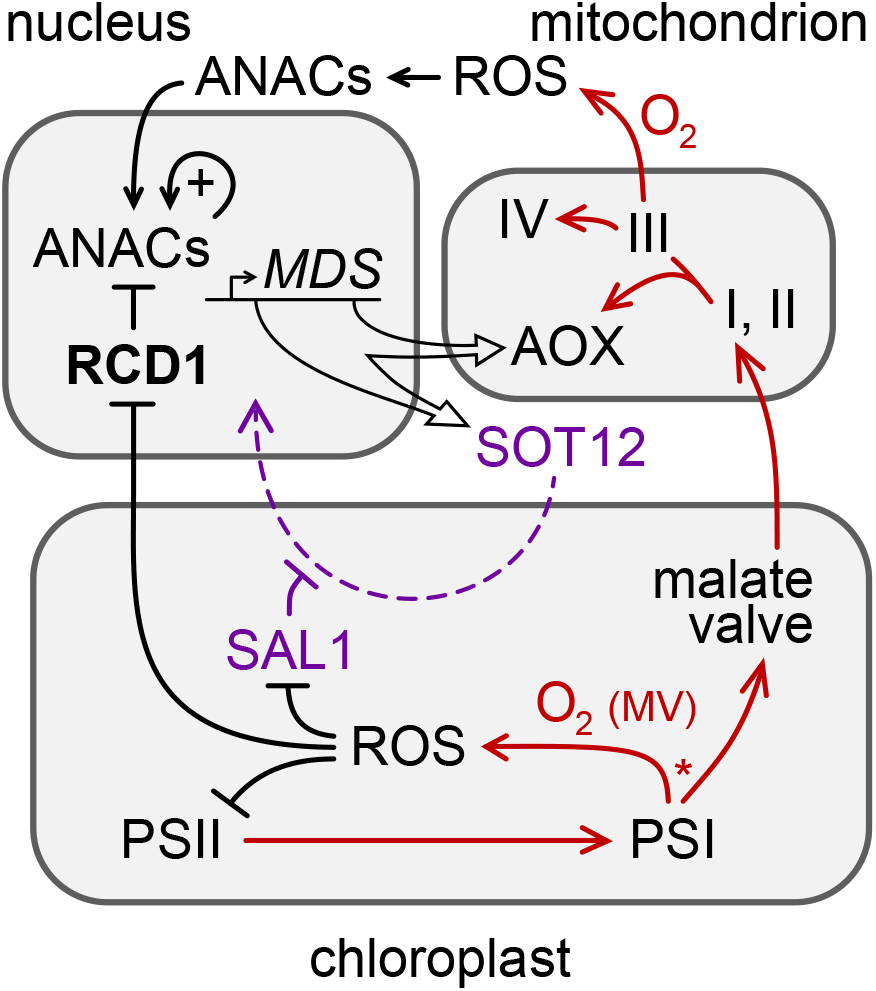
The role of RCD1 in organelle signaling and energy metabolism. RCD1 is the direct suppressor of ANAC transcription factors that is itself subject to redox regulation. Chloroplastic ROS affect RCD1 protein redox state and abundance. Inactivation of RCD1 leads to induction of ANAC-controlled MDS regulon. Expression of MDS genes is feedback-regulated *via* the PAP retrograde signaling (purple). Resulting activation of mitochondrial AOXs and other MDS components affects electron flows (red) and ROS signaling in mitochondria and in chloroplasts. Putative competition of AOX-directed electron transfer with the formation of ROS at PSI is labeled with an asterisk.

Another MDS gene with more abundant mRNA levels in the *rcd1* mutant encodes sulfotransferase SOT12, an enzyme involved in PAP metabolism (Klein and Papenbrock 2004). Accordingly, the SOT12 protein level was significantly increased in the *rcd1* mutant (Fig. 4). Accumulation of SOT12 and similarities between transcript profiles of RCD1- and PAP-regulated genes suggest that PAP signaling is active in *rcd1* mutant. Unbalancing this signaling by elimination of SAL1 leads to severe developmental defects, as evidenced by the stunted phenotype of the *rcd1 sal1* double mutant. Thus, the RCD1 and the PAP signaling pathways appear to be overlapping and somewhat complementary, but the exact molecular mechanisms remain to be explored.

MDS genes represent only a fraction of genes showing differential regulation in *rcd1* (Fig. S3). This likely reflects the fact that RCD1 interacts with many other protein partners in addition to ANACs. The C-terminal RST domain of RCD1 was shown to interact with transcription factors of DREB, PIF, ANAC, Rap2.4 and other families (Jaspers et al. 2009, Vainonen et al. 2012, Hiltscher et al. 2014). Structural analyses of various transcription factors interacting with RCD1 revealed little homology between their RCD1-interacting sequences (O’Shea et al. 2017). Accordingly, the C-terminal domain of RCD1 was proposed to be able to interact with proteins that have different structures. Interaction with ANAC013 significantly changes the structure of this domain in a way that indicates specificity in the interaction (Fig. 5C). Flexible structure of the C-terminal domain of RCD1 probably determines the specificity and ability of RCD1 to interact with different transcription factors, which makes RCD1 a hub in the crosstalk of organellar signaling with hormonal, photoreceptor, immune and other pathways.

The changing environment requires a plant to continuously readjust its energy metabolism and ROS processing. On the one hand, this happens because of abiotic stress factors such as changing light intensity or temperature. For example, a sunlight fleck on a shade-adapted leaf can instantly alter excitation pressure on Photosystems by two orders of magnitude (Allahverdiyeva et al., 2015). On the other hand, chloroplasts and mitochondria are implicated in plant immune reactions to pathogens, contributing to decisive checkpoints including the PCD (Shapiguzov et al., 2012, Petrov et al., 2015, Wu et al. 2015, Van Aken and Pogson 2017, Zhao et al. 2018). In both scenarios, perturbations of organellar ETCs are associated with increased production of ROS, however, the physiological outcomes of the two situations are opposite: acclimation in one case and cell death in the other.

The existence of molecular mechanisms that unambiguously differentiate one type of response from the other has been previously suggested (Trotta et al., 2014, Sowden et al., 2017, Van Aken and Pogson 2017). The ANAC017 transcription factor and MDS genes including *AOX1a*, as well as PAP signaling were proposed as organelle-related components counteracting PCD during abiotic stress (Van Aken and Pogson 2017). This suggests RCD1 to be involved in the regulation of the above cell fate checkpoint. Accordingly, the *rcd1* mutant is characterized by resistance to a number of abiotic stress treatments (Ahlfors et al., 2004, Fujibe et al. 2004, Jaspers et al. 2009).

Interestingly, in contrast to its resistance to abiotic stress, *rcd1* is more sensitive to treatments related to biotic stress. The *rcd1* mutant was originally identified in a forward genetic screen for sensitivity to ozone (Overmyer et al. 2000). Ozone decomposes in the plant cell wall to ROS mimicking formation of ROS by respiratory burst oxidases (RBOHs) in the course of the hypersensitive response in innate immunity (Joo et al., 2005, Vainonen and Kangasjarvi 2015). The opposing roles of RCD1 in the cell fate may be related to its interaction with diverse transcription factor partners and/ or different regulation of its stability and abundance. Indeed, transcriptomic analyses showed that under standard growth conditions, a cluster of genes associated with defense against pathogens had decreased expression in *rcd1* (Brosché et al. 2014), and no ANAC013/ ANAC017 cis-element motif is associated with these genes (Fig. S3). Another possible factor determining varying roles of RCD1 in the cell fate is differential regulation of RCD1 protein function by ROS/ redox signals emitted by different subcellular compartments. The sensitivity of RCD1 to chloroplastic ROS (Fig. 6) can be interpreted as negative regulation of the pro-PCD component. This inactivation can occur in environmental situations that require physiological adaptation rather than PCD. For example, abrupt increase in light intensity can cause excessive electron flow in photosynthetic ETC and overproduction of reducing power. The resulting deficiency of PSI electron acceptors can lead to increased production of chloroplastic ROS, which *via* retrograde signaling influence RCD1 stability and/ or redox status, inhibiting its activity and thus affecting adjustments in nuclear gene expression (Fig. 7). Among other processes, RCD1-mediated suppression of ANAC013/ ANAC017 transcription factors is released, leading to induction of the MDS regulon. The consequent expression of AOXs together with increased chloroplast-to-mitochondrial electron transfer provides electron sink for photosynthesis, which suppresses chloroplast ROS production and contributes to the plant’s survival under changing environment (Fig. 7).

## Materials and methods

### Plants and mutants

*Arabidopsis thaliana* adult plants were grown on soil (peat : vermiculite = 1:1) under white luminescent light (220-250 μmol m^−2^ s^−1^) at a 12-hour photoperiod. Seedlings were grown for 14 days on 1 x MS basal medium (Sigma-Aldrich) with 0.5 % Phytagel (Sigma-Aldrich). Arabidopsis *rcd1-4* mutant (GK-229D11), *rcd1-1* (Overmyer et al. 2000), *aox1a* (SAIL_030_D08), *AOX1a-OE* (Umbach et al. 2005), *ptox* (Wetzel et al., 1994), *anac017* (SALK_022174), and *sal1-8* (Wilson et al., 2009) mutants are of Col-0 background; *pgr5* mutant is of *gl1* background (Munekage et al., 2002). ANAC013-GFP line is described in (De Clercq et al. 2013), RCD1-HA line < in (Jaspers et al. 2009). RCD1-3xVenus, rcd1Δ7Cys-HA lines are described in *Cloning.*

### Cloning

To create the RCD1-3xVenus construct, RCD1 cDNA was fused to the *UBIQUITIN10* promoter region and to the C-terminal triple Venus YFP tag in a MultiSite Gateway reaction as described in (Siligato et al., 2016). To create the rcd1Δ7Cys-HA construct, cysteine residues were mutated to alanines by sequential PCR-based mutagenesis of the genomic sequence of RCD1 in the pDONR/Zeo vector. The resulting construct was transferred to the pGWB13 binary vector by a Gateway reaction and introduced to the *rcd1-4* mutant by floral dipping. Homozygous single insertion lines were obtained. For HEK293T cell experiments codon-optimized N-terminal 3xHA-fusion of RCD1 and C-terminal 3xmyc-fusion of ANAC013 were cloned into pcDNA3.1(+). Full-length ANAC017 was cloned pcDNA3.1(-) in the Xho I/ Hind III sites, the double myc tag was introduced in the reverse primer sequence.

### Inhibitor treatments

For photoinhibition studies, leaf discs were let floating on Milli-Q water solution supplemented with 0.05 % Tween 20 (Sigma-Aldrich). Final dilution of AA and myx was 2.5 μM each, of SHAM – 2 mM. Stock solutions of these chemicals were prepared in DMSO, equal volumes of DMSO were added to control samples. Pretreatment with chemicals was carried out in the darkness, overnight for MV, AA and myx, 1 hour for SHAM. For chemical treatment in seedlings grown on MS plates, 5 mL of Milli-Q water with or without 50 μM MV were poured in 9-cm plates at the end of the light period.

### DAB staining

Plant rosettes were stained with 3,3’-diaminobenzidine (DAB) essentially as described in (Daudi et al., 2012) [Daudi, A. and O’Brien, J. A. (2012). Detection of Hydrogen Peroxide by DAB Staining in Arabidopsis Leaves. Bio-protocol 2(18): e263. DOI: 10.21769/BioProtoc.263.]. After vacuum infiltration of DAB-staining solution in the darkness, rosettes were exposed to light (180 μmol m^−2^ s^−1^) for 20 min to induce production of chloroplastic ROS and then immediately transferred to the bleaching solution.

### Spectroscopic measurements of photosynthesis

Chlorophyll fluorescence was measured by MAXI Imaging PAM (Walz, Germany). PSII photoinhibition protocol consisted of repetitive 1-hour periods of blue actinic light (450 nm, 80 μmol m^−2^ s^−1^) each followed by a 20-min dark adaptation, then Fo and Fm measurement. PSII photochemical yield was calculated as Fv/Fm = (Fm-Fo)/Fm. To plot raw chlorophyll fluorescence kinetics under light (Fs) against time, the reads were normalized to dark-adapted Fo. The assays were performed in 96-well plates. In each assay leaf discs from at least 4 individual plants were analyzed. Each assay was reproduced at least three times.

PSI (P700) oxidation was measured by DUAL-PAM-100 (Walz, Germany) as described (Tiwari et al., 2016). Leaves were pre-treated in 1 μM MV for 4 hours, then shifted to light (160 μmol m^−2^ s^−1^) for indicated time. Oxidation of P700 was induced by PSI-specific far red light (FR, 720 nm). To determine fully oxidized P700 (Pm), a saturating pulse of actinic light was applied under continuous background of FR, followed by switching off both the actinic and FR light. The kinetics of P700+ reduction by intersystem electron transfer pool and re-oxidation by FR was determined by using a multiple turnover (MT) saturating flash of PSII light (635 nm) in the background of continuous FR.

### Isolation, separation and detection of proteins and protein complexes

Thylakoids were isolated as described in (Jarvi et al., 2016) and leaf total protein as described in (Kangasjarvi et al., 2008). Chlorophyll content was determined according to (Porra et al., 1989) and protein content according to (Lowry et al., 1951).

For separation of proteins SDS-PAGE (12 % polyacrylamide) was used (Laemmli 1970). For thylakoid proteins the gel was complemented with 6 M urea. To separate thylakoid membrane protein complexes, isolated thylakoids were solubilized with n-dodecyl β-D-maltoside (Sigma-Aldrich) and separated in BN-PAGE (5-12.5 % polyacrylamide) as described by (Jarvi et al. 2016).

For immunoblotting of total plant extracts the plant material was frozen immediately after treatments in liquid nitrogen and ground. Total proteins were extracted in SDS extraction buffer [50 mM Tris, pH 7.8, 2 % SDS, 1 x protease inhibitor cocktail (Sigma-Aldrich), 2 mg/ mL NaF] for 20 min at 37 °C and centrifuged at 18 000 x g for 10 min. Supernatants were normalized for protein concentration and resolved by SDS-PAGE. After electrophoresis, proteins were electroblotted to PVDF membrane and immunoblotted with specific antibodies.

### Analysis of protein thiol redox state by mobility shift assays

Thiol redox state of 2-CPs in detached Col-0 and *rcd1* leaves adapted to darkness or light (3 hours of 160 μmol m^−2^ s^−1^), was determined by alkylating free thiols in TCA-precipitated proteins with 50 mM N-ethylmaleimide, reducing *in vivo* disulfides with 100 mM DTT and then alkylating the newly reduced thiols with 10 mM methoxypolyethylene glycol maleimide of molecular weight 5 kDa (MMPEG) (Sigma-Aldrich), as described in (Nikkanen et al., 2016). Proteins were then separated by SDS-PAGE and immunoblotted with a 2-CP-specific antibody.

### Preparation of crude mitochondria

Crude mitochondria were isolated from Arabidopsis rosette leaves as described in (Keech et al., 2005).

### *Measurements of AOX capacity* in vivo

Seedling respiration and AOX capacity were assessed by measuring O_2_ consumption in the darkness using a Clark electrode as described in (Schwarzländer et al., 2009).

### Non-aqueous fractionation (NAF)

Leaves of Arabidopsis plants were harvested and snap-frozen in liquid nitrogen. 4 grams of fresh weight of frozen plant material was ground to a fine powder using a mixer mill (Retsch), transferred to Falcon tubes and freeze-dried at 0.02 bar for 5 days in a lyophilizer which had been pre-cooled to −40 °C. The NAF-fractionation procedure was performed as described in (Krueger et al., 2011, Arrivault et al., 2014, Krueger et al., 2014) except that the gradient volume, composed of the solvents tetrachlorethylene (C2Cl4)/ heptane (C7H16), was reduced from 30 mL to 25 mL but with a same linear density. Leaf powder was resuspended in 20 mL C2Cl4/ C7H16 mixture 66:34 (v/v; density p = 1.3 g cm^−3^), and sonicated for 2 min, with 6 × 10 cycles at 65 % power. The sonicated suspension was filtered through a nylon net (20 μm pore size). The net was washed with 30 mL of heptane. The suspension was centrifuged for 10 min at 3 200 x g and 4 °C and the pellet was resuspended in 5 mL C2Cl4/ C7H16 mixture 66:34. The gradient was formed in 38 mL polyallomer centrifugation tube using a peristaltic gradient pump (BioRad) generating a linear gradient from 70 % solvent A (C2Cl4/ C7H16 mixture 66:34) to 100 % solvent B (100 % C2Cl4) with a flow rate of 1.15 mL min^−1^, resulting in a density gradient from 1.43 g cm^−3^ to 1.62 g cm^−3^. 5 mL suspension containing the sample was loaded on top of the gradient and centrifuged for 55 min at 5 000 x g and 4 °C using a swing-out rotor with acceleration and deceleration of 3:3 (brakes off). Each of the compartment-enriched fractions (F1 to F8) were transferred carefully from the top of the gradient into a 50 mL Falcon tube and filled up with heptane to a volume of 20 mL and centrifuged at 3 200 x g for 10 min. The pellet was resuspended in 6 mL of heptane and subsequently divided into 6 aliquots of equal volume (950 μL). The pellets had been dried in a vacuum concentrator without heating and stored at −80 °C until further use. Subcellular compartmentation of markers or the metabolites of our interest was calculated by BestFit method as described in (Krueger et al. 2011, Krueger et al. 2014). Percentage values (% of the total found in all fractions) of markers and metabolites have been used to make the linear regressions for subcellular compartments using BestFit.

### Marker measurements for non-aqueous fractionation

Before enzyme and metabolite measurements dried pellets were homogenized in the corresponding extraction buffer by the addition of one steel ball (2-mm diameter) to each sample and shaking at 25 Hz for 1 min in a mixer mill. Enzyme extracts were prepared as described in (Gibon et al., 2004) with some modifications. The extraction buffer contained 50 mM HEPES/ KOH, pH 7.5, 10 mM MgCl2, 1 mM EDTA, 1 mM EGTA, 1 mM benzamidine, 1 mM ε-aminocapronic acid, 0.25 % (w/v) BSA, 20 μM leupeptin, 0.5 mM DTT, 1 mM phenylmethylsulfonyl fluoride (PMSF), 1 % (v/v) Triton X-100, 20 % glycerol. The extract was centrifuged (14 000 rpm for 10 min at 4°C) and the supernatant was used directly for the enzymatic assays. The activities of adenosine diphosphate glucose pyrophosphorylase (AGPase) and phosphoenolpyruvate carboxylase (PEPC) were determined as described in (Gibon et al. 2004) but without using the robot-based platform as described in the paper. Chlorophyll was extracted twice with 80 % (v/v) and once with 50 % (v/v) hot ethanol/ 10 mM HEPES, pH 7.0 followed by 30-min incubation at 80 °C and determined as described in (Cross et al., 2006). Nitrate was measured by the enzymatic reaction as described in (Cross et al. 2006).

### Incubation of Arabidopsis leaf discs with [U-^14^C] glucose

For the light experiment, leaf discs were incubated under light in 5 mL 10 mM MES-KOH, pH 6.5, containing 1.85 MBq/ mmol [U-^14^C] glucose (Hartmann Analytic) in a final concentration of 2 mM. In the dark experiment, leaf discs were incubated under green light for 150 min. Leaf discs were placed in a sieve, washed several times in double-distilled water, frozen in liquid nitrogen, and stored at −80 °C until further analysis. All incubations were performed in sealed flasks under green light and shaken at 100 rpm. The evolved ^14^CO_2_ was collected in 0.5 mL of 10 % (w/v) KOH.

### Fractionation of ^14^C-labeled tissue extracts and measurement of metabolic fluxes

Extraction and fractionation were performed according to (Obata et al., 2017). Frozen leaf discs were extracted with 80 % (v/v) ethanol at 80 °C (1 mL per sample) and re-extracted in two subsequent steps with 50 % (v/v) ethanol (1 mL per sample for each step), and the combined supernatants were dried under an air stream at 35 °C and resuspended in 1 mL of water (Fernie et al., 2001). The soluble fraction was subsequently separated into neutral, anionic, and basic fractions by ion-exchange chromatography; the neutral fraction (2.5 mL) was freeze-dried, resuspended in 100 μL water, and further analyzed by enzymatic digestion followed by a second ion-exchange chromatography step (Carrari et al., 2006). To measure phosphate esters, samples (250 μL) of the soluble fraction were incubated in 50 μL of 10 mM MES-KOH, pH 6.0, with or without 1 unit of potato acid phosphatase (grade II; Boehringer Mannheim) for 3 hours at 37 °C, boiled for 2 min, and analyzed by ion-exchange chromatography (Fernie et al. 2001). The insoluble material left after ethanol extraction was homogenized, resuspended in 1 mL of water, and counted for starch (Fernie et al. 2001). Fluxes were calculated as described following the assumptions detailed by Geigenberger *et al* (Geigenberger et al., 1997, Geigenberger et al., 2000).

### Metabolite extraction

Primary metabolites were analyzed with GC-MS according to (Roessner et al., 2000). GC-MS analysis was executed from the plant extracts of eight biological replicates (pooled samples). Plant material was homogenized in a Qiagen Tissuelyser II bead mill (Qiagen, Germany) with 1-1.5 mm Retsch glass beads. Soluble metabolites were extracted from plant material in two steps, first with 1 mL of 100 % methanol (Merck) and second with 1 mL of 80 % (v/v) aqueous methanol. During the first extraction step, 5 μL of internal standard solution (0.2 mg mL^−1^ of benzoic-ds acid, 0.1 mg mL^−1^ of glycerol-d_5_, 0.2 mg mL^−1^ of 4-methylumbelliferone in methanol) was added to each sample. During both extraction steps, the samples were vortexed for 30 min and centrifuged for 5 min at 13 000 rpm (13 500 × g) at 4 °C. The supernatants were then combined for metabolite analysis. The extracts (2 mL) were dried in a vacuum concentrator (MiVac Duo, Genevac Ltd, Ipswich, UK), the vials were degassed with nitrogen and stored at −80 °C prior to derivatization and GC-MS analysis.

Dried extracts were re-suspended in 500 μL of methanol. Aliquot of 200 μL was transferred to a vial and dried in a vacuum. The samples were derivatized with 40 μL of methoxyamine hydrochloride (MAHC, Sigma-Aldrich) (20 mg mL^−1^) in pyridine (Sigma-Aldrich) for 90 min at 30 °C at 150 rpm, and with 80 μL N-methyl-N-(trimethylsilyl) trifluoroacetamide with 1 % trimethylchlorosilane (MSTFA with 1 % TMCS, Thermo Scientific) for 120 min at 37 °C at 150 rpm. Alkane series (10 μL, C10–C40, Supelco) in hexane (Sigma-Aldrich) and 100 μL of hexane was added to each sample before GC-MS analysis.

### Metabolite analysis by gas chromatography-mass spectrometry

The GC-MS system consisted of Agilent 7890A gas chromatograph with 7000 Triple quadrupole mass spectrometer and GC PAL autosampler and injector (CTC Analytics). Splitless injection (1 μL) was employed using a deactivated single tapered splitless liner with glass wool (Topaz, 4 mm ID, Restek). Helium flow in the column (Agilent HP-5MS Ultra Inert, length 30 m, 0.25 mm ID, 0.25 μm film thickness combined with Agilent Ultimate Plus deactivated fused silica, length 5 m, 0.25 mm ID) was 1.2 mL min^−1^ and purge flow at 0.60 min was 50 mL min^−1^. The injection temperature was set to 270 °C, MS interface 180 °C, source 230 °C and quadrupole 150 °C. The oven temperature program was as follows: 2 min at 50 °C, followed by a 7 °C min^−1^ ramp to 260 °C, 15 °C min^−1^ ramp to 325 °C, 4 min at 325 °C and post-run at 50 °C for 4.5 min. Mass spectra were collected with a scan range of 55-550 *m/z.*

Metabolite Detector (versions 2.06 beta and 2.2N) (Hiller et al., 2009) and AMDIS (version 2.68, NIST) were used for deconvolution, component detection and quantification. Malate and fumarate levels were calculated as the peak area of the metabolite normalized with the peak area of the internal standard, glycerol-d_8_, and the fresh weight of the sample.

### Measurements of NADPH-MDH activity

From light-adapted plants grown for 5 weeks at a 8-hour day photoperiod, total extracts were prepared as for non-aqueous fractionation in the extraction buffer supplemented with 250 μM DTT. In microplates 5 μL of the extract (diluted x 500) was mixed with 20 μL of activation buffer (0.1 M Tricine-KOH, pH 8.0, 180 mM KCl, 0.5 % Triton X-100, with or without DTT 150 mM) and incubated for 2 hours at room temperature. Then assay mix was added consisting of 20 μL of assay buffer (0.5 M Tricine-KOH, pH 8.0, 0.25 % Triton X-100, 0.5 mM EDTA), 9 μL of water, and 1 μL of 50 mM NADPH (prepared in 50 mM NaOH), after which 45 μL of 2.5 mM oxaloacetate or water control was added. The reaction was mixed, and light absorbance at 340-nm wavelength was measured at 25 °C.

### *Analysis of* rcd1 *misregulated genes in microarray experiments related to chloroplast or mitochondrial dysfunction*

Genes with misregulated expression in *rcd1* were selected from our previous microarray datasets (Brosché et al. 2014) with the cutoff, absolute value of logFC < 0.5. These genes were subsequently clustered with the *rcd1* gene expression dataset together with various Affymetrix datasets related to chloroplast or mitochondrial dysfunction from the public domain using bootstrapped Bayesian hierarchical clustering as described in (Wrzaczek et al., 2010). Affymetrix raw data (.cel files) were normalized with Robust Multi-array Average normalization, and manually annotated to control and treatment conditions, or mutant *versus* wild type.

Affymetrix ATH1-121501 data were from the following sources: Gene Expression Omnibus https://www.ncbi.nlm.nih.gov/geo/, AA 3 hours (in figures labelled as experiment 1), GSE57140 (Ivanova et al., 2014); AA and H_2_O_2_, 3 hour treatments (in figures labelled as experiment 2), GSE41136 (Ng et al. 2013); MV 3 hours, GSE41963 (Sharma et al., 2013); *mterf6-1,* GSE75824 (Leister and Kleine 2016); *prors1-2,* GSE54573 (Leister et al., 2014); H_2_O_2_ 30 min, GSE43551 (Gutiérrez et al., 2014); high light 1 hour (in figures labelled as experiment 1), GSE46107 (Van Aken et al., 2013); high light 30 min in cell culture, GSE22671 (González-Pérez et al., 2011); high light 3 hours (in figures labelled as experiment 2), GSE7743 (Kleine et al., 2007); oligomycin 1 and 4 hours, GSE38965 (Geisler et al., 2012); norflurazon – 5 day-old seedlings grown on plates with norflurazon, GSE12887 (Koussevitzky et al., 2007); *msh1 recA3* double mutant, GSE19603 (Shedge et al., 2010). AtGenExpress oxidative time series, MV 12 and 24 hours, http://www.arabidopsis.org/servlets/TairObject?type=expressionset&id=1007966941. ArrayExpress, https://www.ebi.ac.uk/arrayexpress/: rotenone, 3 and 12 hours, E-MEXP-1797 (Garmier et al., 2008); *alx8* and *fry1,* E-MEXP-1495 (Wilson et al. 2009); *ndufs4,* E-MEXP-1967 (Meyer et al., 2009).

### Identification of interacting proteins using IP/MS-MS

Immunoprecipitation experiments were performed in three biological replicates as described previously (De Rybel et al., 2013), using 3 g of rosette leaves from p35S:ANAC013-GFP and 2.5 g of rosette leaves from pUBI10:RCD1-3xVenus transgenic lines. Interacting proteins were isolated by applying total protein extracts to αGFP-coupled magnetic beads (Milteny Biotech). Three replicates of p35S:ANAC013-GFP or pUBI10:RCD1-3xVenus were compared to three replicates of Col-0 controls. Tandem mass spectrometry (MS) and statistical analysis using MaxQuant and Perseus software was performed as described previously (Wendrich et al., 2017).

### HEK293T human embryonic kidney cell culture and transfection

HEK293T cells were maintained at 37 °C and 5 % CO_2_ in Dulbecco’s Modified Eagle’s Medium F12-HAM, supplemented with 10 % foetal bovine serum, 15 mM HEPES, and 1 % penicillin/ streptomycin. Cells were transiently transfected using GeneJuice (Novagen) according to the manufacturer’s instructions.

For co-immunoprecipitation experiments, HEK293T cells were co-transfected with plasmids encoding HA-RCD1 and ANAC013-myc or ANAC017-myc. Forty hours after transfection, cells were lysed in TNE buffer [50 mM Tris-HCl, pH 7.4, 150 mM NaCl, 5 mM EDTA, 1 % Triton X-100, 1 x protease inhibitor cocktail, 50 μM proteasome inhibitor MG132 (Sigma-Aldrich)]. After incubation for 2 hours at 4 °C, lysates were cleared by centrifugation at 18 000 x g for 10 min at 4 °C. For co-immunoprecipitations, cleared cell lysates were incubated with either αHA or αmyc antibody immobilized on agarose beads overnight at 4 °C. Beads were washed six times with the lysis buffer. The bound proteins were dissolved in SDS sample buffer, resolved by SDS-PAGE, and immunoblotted with the specified antibodies.

### Protein expression and purification

The C-terminal domain of RCD1 for NMR study was expressed as GST-fusion protein in *E.coli* BL21 (DE3) Codon Plus strain and purified using GSH-Sepharose beads (GE Healthcare) according to the manufacturer’s instruction. Cleavage of GST tag was performed with thrombin (GE Healthcare, 80 units per mL of beads) for 4 hours at room temperature and the C-terminal domain of RCD1 was eluted from the beads with PBS buffer (137 mM NaCl, 2.7 mM KCl, 10 mM Na2HPO4, 1.8 mM KH2PO4, pH 7.4). The protein was further purified by gel filtration with HiLoad 16/600 Superdex 75 column (GE Healthcare) equilibrated with 20 mM sodium phosphate buffer, pH 6.4, 50 mM NaCl at 4 °C.

### Peptide synthesis

ANAC013 peptides of > 98 % purity for surface plasmon resonance and NMR analysis were purchased from Genecust, dissolved in water to 5 mM final concentration and stored at −80 °C before analyses.

### Surface plasmon resonance

The C-terminal domain of RCD1 was covalently coupled to a Biacore CM5 sensor chip *via* amino-groups. 500 nM of ANAC013 peptides were then profiled at a flow rate of 30 μL min^−1^ for 300 s, followed by 600 s flow of wash buffer. After analysis in BiaEvalution (Biacore) software, the normalized resonance units were plotted over time with the assumption of one-to-one binding.

### NMR spectroscopy

NMR sample production and chemical shift assignment have been described in (Tossavainen et al. 2017). A Bruker Avance III HD 800 MHz spectrometer equipped with a TCI ^1^H/ ^13^C/ ^15^N cryoprobe was used to acquire spectra for structure determination of RCD1^468-589^. Peaks were manually picked from three NOE spectra, a ^1^H, ^15^N NOESY-HSQC and ^1^H, ^13^C NOESY-HSQC spectra for the aliphatic and aromatic ^13^C regions. CYANA 2.1 (Lopez-Mendez and Guntert 2006) automatic NOE peak assignment – structure calculation routine was used to generate 300 structures from which 30 were further refined in explicit water with AMBER 16 (Case et al., 2005). Assignments of three NOE peaks were kept fixed using the KEEP subroutine in CYANA. These NOE peaks restrained distances between the side chains of W507 and M508 and adjacent helices 1 and 4, respectively. Fifteen lowest AMBER energy structures were chosen to represent of RCD1^468-589^ structure in solution.

Peptide binding experiment was carried out by preparing a sample containing of RCD1^468-589^ and ANAC013^235-284^ peptide in an approximately 1:2 concentration ratio, and recording a ^1^H, ^15^N HSQC spectrum. Amide peak positions were compared with those of the free RCD1^468-589^.

## Acknowledgements

We thank Dr. Olga Blokhina, Dr. Bernadette Gehl and Manuel Saornil for the help in studies of mitochondria, Katariina Vuorinen for genotyping *rcd1-1 anac017,* Richard Gossens for critical comments on the manuscript. Prof. F. J. Cejudo (Institute of Plant Biochemistry, University of Sevilla) for the α2-CP antibody. We acknowledge CSC – IT Center for Science, Finland, for computational resources. This work was supported by the University of Helsinki (JK); the Academy of Finland Centre of Excellence programs (2006-11; JK and 2014-19; JK, EMA) and Research Grant (Decision 250336; JK); Academy of Finland fellowships (Decisions 275632, 283139 and 312498; MW) Academy of Finland Research Grant (Decision 288235; PP); by The Research Foundation – Flanders (FWO; Odysseus II G0D0515N and Post-doc grant 12D1815N; BW and BDR); PlantaSYST project by the European Union’s Horizon 2020 research and innovation programme (SGA-CSA No 664621 and No 739582 under FPA No. 664620; SA and ARF); the Research Foundation-Flanders (Excellence of Science project no 30829584; FVB).

## Author contributions

J.K., A.S., M.W., S.J., J.P.V., M.B., A.R.F., E.R., E.-M.A., F.B., and P.P. designed the research. J.P.V., K.H., B.W., K.V.D.K., B.D.R., F.B., and M.W. carried out *in vivo* pulldown experiments of RCD1 and ANACs. A.S., J.P.V., S.J., A.T., L.N., E.T., E.R., and E.-M.A. performed studies of photosynthesis and respiration. A.R.F., F.A., S.A., N.S., A.S., and M.K. conducted metabolomic studies. J.P.V., H.T., M.H., and P.P. carried out structural analyses of RCD1. J.S. and M.B. performed bioinformatic studies. A.S., J.P.V., M.W., J.K.-W., M.B., A.R.F., and J.K. wrote the manuscript.

## Supplementary Information

**Supplementary Table 1. Metabolic analyses.**

Distribution of radioactive label was analyzed after feeding plants with ^14^C-labeled glucose, and deduced metabolic fluxes in light- and dark-adapted Col-0, *rcd1, rcd1 aox1a,* and *aox1a* plants.

**Supplementary Table 2. *In vivo* interaction partners of ANAC013.**

From Arabidopsis line expressing ANAC013-GFP, ANAC013-GFP and associated proteins were purified with αGFP antibody and identified by mass spectrometry. Identified proteins (Perseus analysis, ANAC013) and mapped peptides (peptide IDs) are shown.

**Supplementary Table 3. *In vivo* interaction partners of RCD1.**

From Arabidopsis line expressing RCD1-3xVenus, RCD1-3xVenus and associated proteins were purified with αGFP antibody and identified by mass spectrometry. Identified proteins (Perseus analysis, RCD1) and mapped peptides (peptide IDs) are shown.

**Supplementary Table 4. NMR constraints and structural statistics for the ensemble of the 15 lowest-energy structures of RCD1 RST.**

**Supplementary Table 5. Statistical analyses.**

The statistical analyses made for the datasets presented in the corresponding figures.

---

**Figure S1.**
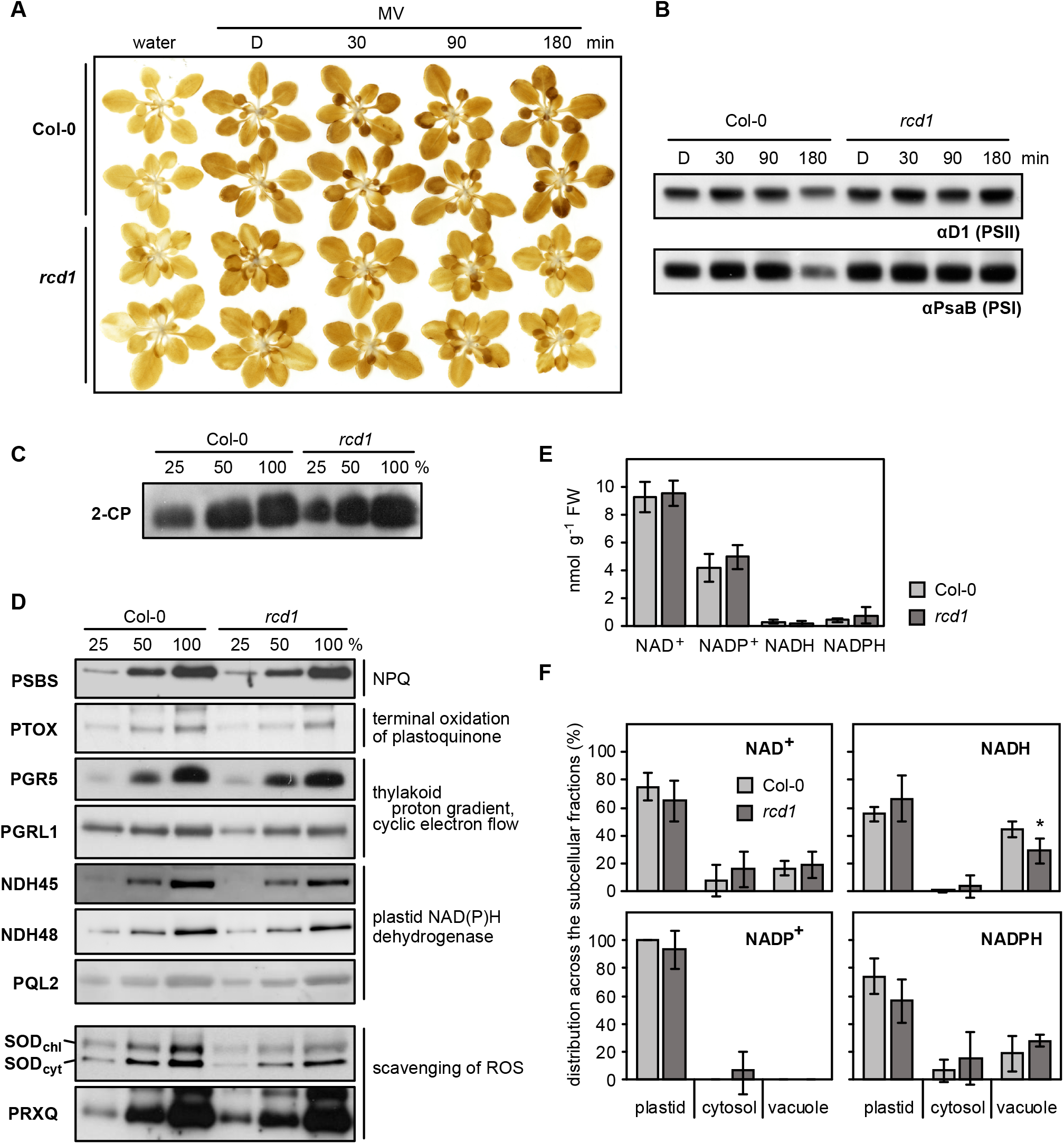
Alterations in *rcd1* chloroplasts. (A) H_2_O_2_ accumulation in rosettes of Col-0 and *rcd1* under MV treatment in the light at indicated time points, revealed by DAB staining (D – dark control). (B) Abundance of PSII (αD1 antibody) and PSI (αPsaB antibody) in protein extracts isolated from MV-treated leaves that were exposed to light for indicated times (D – dark control). Proteins were separated in denaturing SDS-PAGE. (C) Abundance of chloroplastic 2-CP in Col-0 and *rcd1* assessed by separation in SDS-PAGE and immunostaining with the specific antibody. (D) Abundance of proteins involved in different chloroplastic processes assessed by separation in SDS-PAGE and immunostaining with specific antibodies. (E) Abundance of nucleotides NAD+, NADP+, NADH and NADPH in Col-0 and *rcd1* (mean ± SE). (F) Distribution of NAD+/ NADH and NADP+/ NADPH redox couples in various cellular compartments of Col-0 and *rcd1,* assessed by non-aqueous fractionation metabolomics (mean ± SE, an asterisk indicates the value significantly different from that in the corresponding wild type, *P value < 0.05, Student’s t-test).

**Figure S2.**
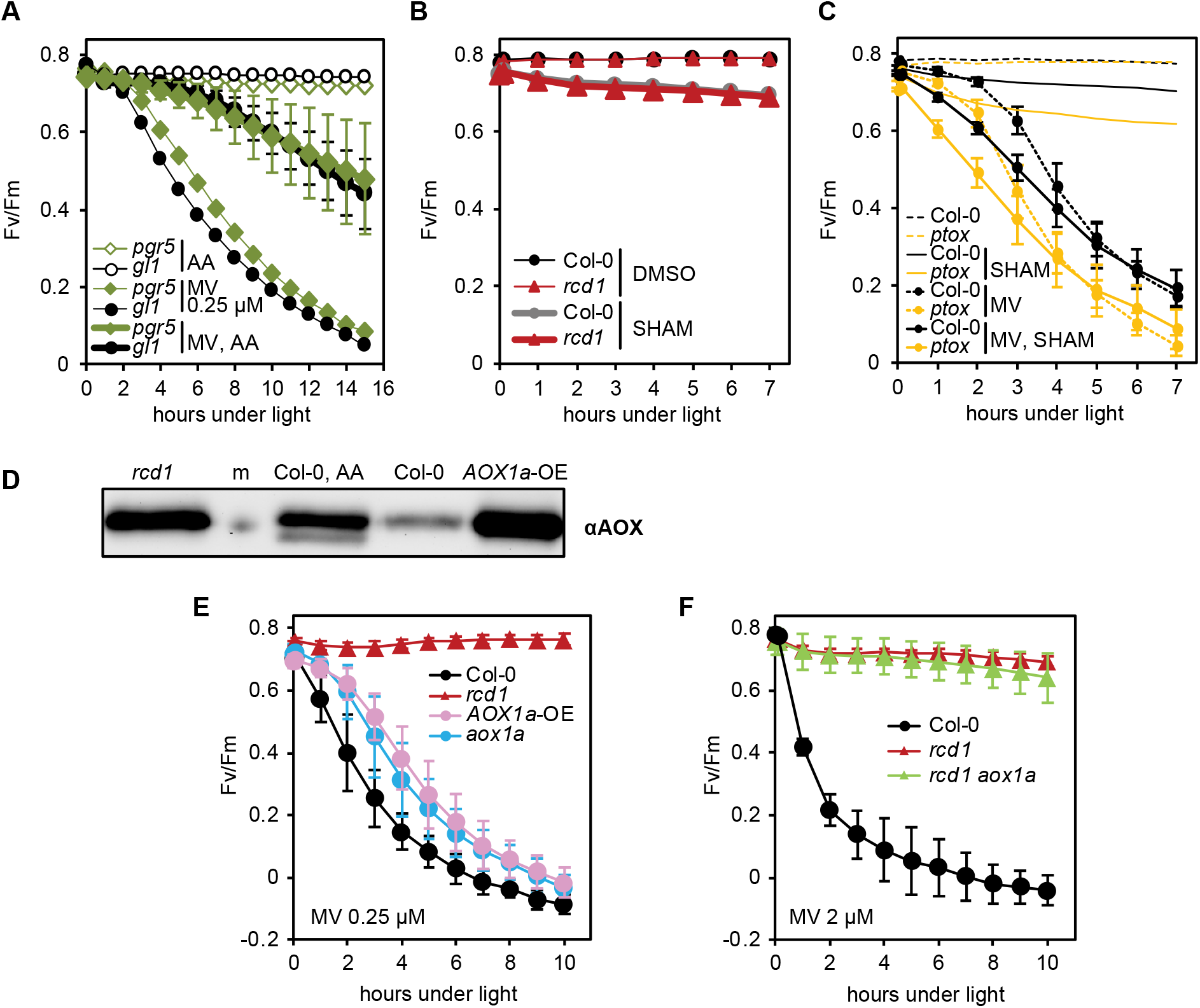
Specificity of inhibitor treatments, effect of SHAM on PSII photoinhibition and irrelevance of AOX1a isoform for MV tolerance. All chlorophyll fluorescence analyses are presented as mean ± SD, for full statistical analysis see Supplementary Table 5. (A) Effect of AA on MV-induced PSII photoinhibition in *pgr5* and in its corresponding wild type *gl1.* No significant difference was observed between the tested lines (Supplementary Table 5). (B) SHAM-induced PSII photoinhibition. No significant difference was detected between the genotypes (Supplementary Table 5). (C) Effect of SHAM on MV-induced PSII photoinhibition in green sectors of leaves of the *ptox* mutant. (D) Abundance of total AOX in the AOX1a-overexpressor line *(AOX1a-OE)* (m – molecular weight marker). (E) MV-induced PSII photoinhibition in the *AOX1a-OE* and *aox1a* lines. No significant difference was observed between *AOX1a-OE* and *aox1a* at any time point of the experiment (Supplementary Table 5). (F) MV-induced PSII photoinhibition in *rcd1 aox1a* double mutant. No significant difference was detected between *rcd1 aox1a* and *rcd1* (Supplementary Table 5).

**Figure S3.**
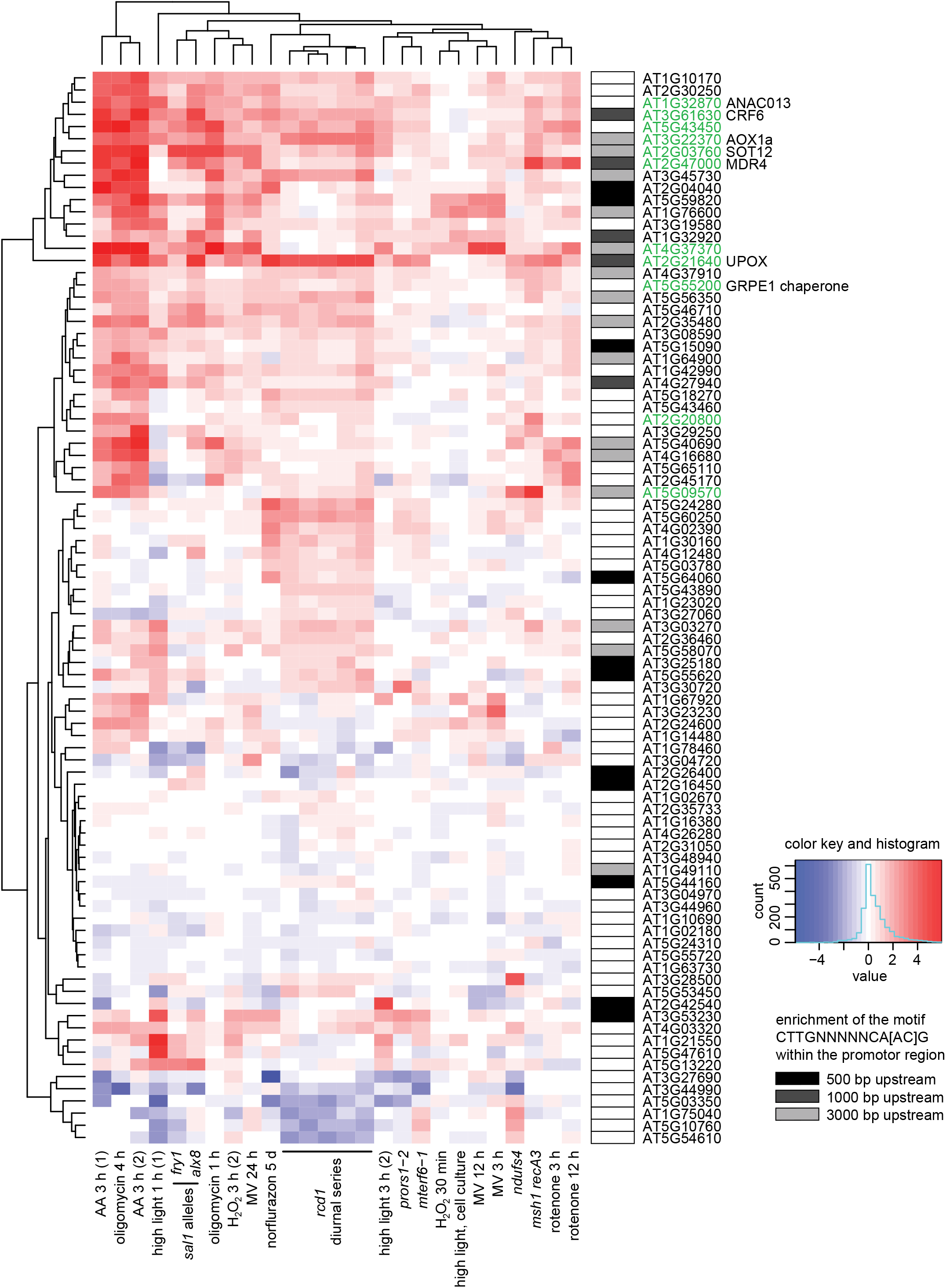
Clustering analysis of genes mis-regulated in *rcd1* (with cutoff of logFC < 0.5) in published gene expression data sets acquired after perturbations of chloroplasts or mitochondria. MDS genes are labeled green. Enrichment of the ANAC013/ ANAC017 cis-element CTTGNNNNNCA[AC]G (De Clercq et al. 2013) in promoter regions is shown by shaded boxes next to the gene names.

**Figure S4.**
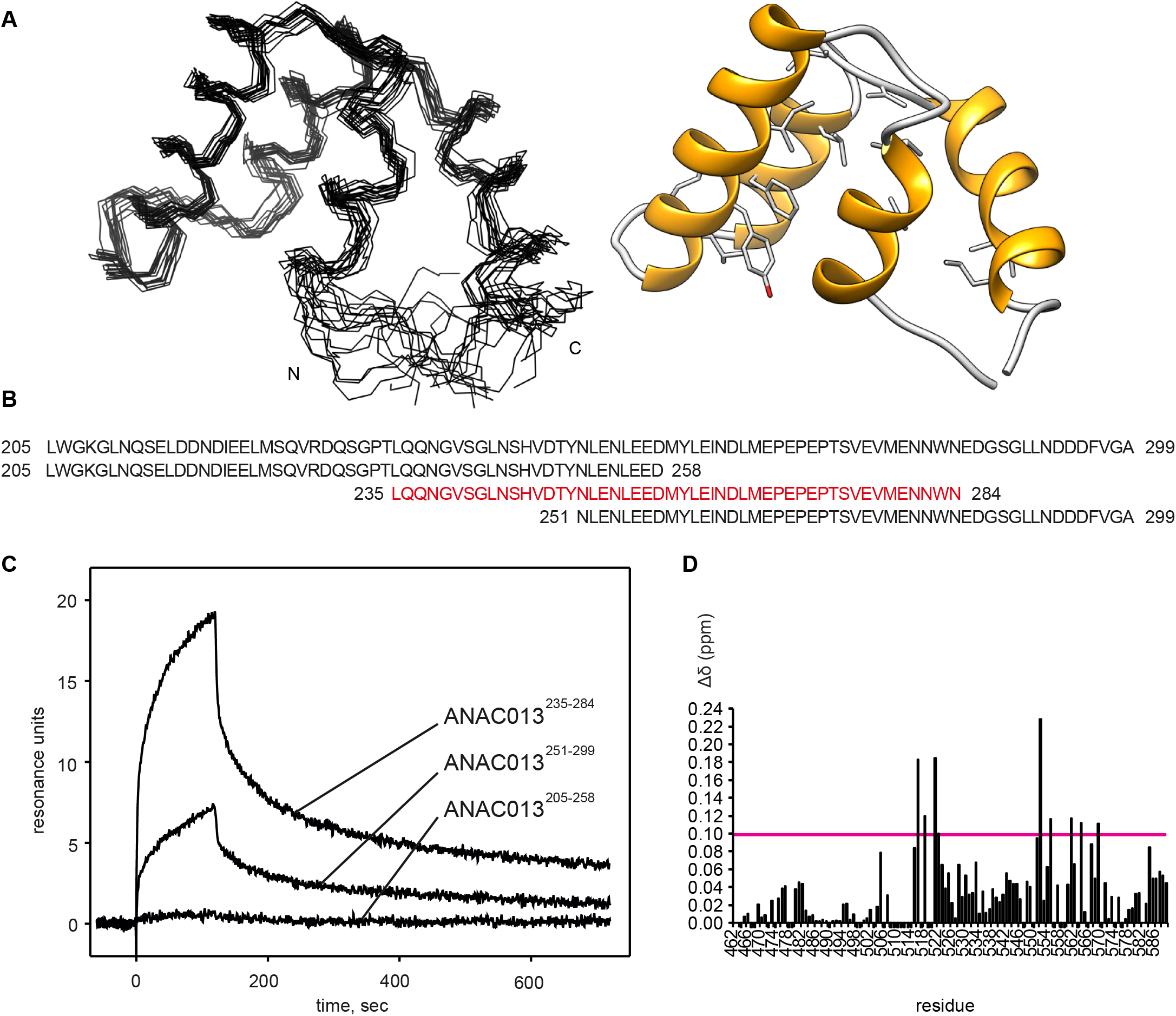
Structure of the RST domain of RCD1 and analysis of ANAC013-derived peptides for interaction with the RST domain. (A) Structure of the C-terminal domain of RCD1 (residues G468-L589) was determined by NMR spectroscopy. The first 38 N-terminal and the last 20 C-terminal are devoid of any persistent structure, hence only the structure of the folded part (residues P506-P570) is shown. The ensemble of 15 lowest-energy structures is on the left and a ribbon representation of the lowest-energy structure on the right. The domain consists of four α-helices, F510-I517, E523-R537, R543-V554 and D556-L566. The structure of the beginning of the first helix is dispersed in the ensemble due to sparseness of distance restraints. This arises from several missing amide chemical shift assignments (Tossavainen et al. 2017) as well as the presence of four proline residues in this region (P503, P506, P509 and P511) which severely hindered distance restraint generation. The many conserved hydrophobic residues (Jaspers et al. 2010), shown in stick representation, form the domain’s hydrophobic core. Mutagenesis experiments identified hydrophobic residues L528/I529 and I563 as critical for RCD1 interaction with DREB2A (Vainonen et al. 2012). I529 and I563 are constituents of the hydrophobic core, and substitution of these residues probably disrupts the core of the RST domain thus abolishing the interaction. (B) According to yeast two-hybrid data (O’Shea et al. 2017), ANAC013 residues 205-299 are responsible for interaction with RCD1. To narrow down the RCD1-interacting domain, we designed three overlapping peptides ANAC013^205-258^, ANAC013^235-284^, ANAC013^251-299^ and tested their binding to RCD1 by surface plasmon resonance. (C) Surface plasmon resonance interaction analysis of three ANAC013-derived peptides with the C-terminal domain of RCD1. The strongest binding was detected for ANAC013 peptide 235-284 (red in panel B), which was further used for the NMR titration experiment with the purified C-terminal domain of RCD1. (D) Histogram depicting the changes in ^1^H and ^15^N chemical shifts in RCD1^468-589^ upon addition of the ANAC013^235-284^ peptide. Changes were quantified according to the ‘minimum chemical shift procedure’, that is, each peak in the free form spectrum was linked to the nearest peak in the bound form spectrum. An arbitrary value −0.005 ppm was assigned to residues for which no data could be retrieved. Residues that exhibited the largest changes (≥ 0.10 ppm) are identified by residue number in the ^1^H, ^15^N HSQC spectrum and highlighted on the RS_TRCD1_ structure in Fig. 5C inset.

**Figure S5.**
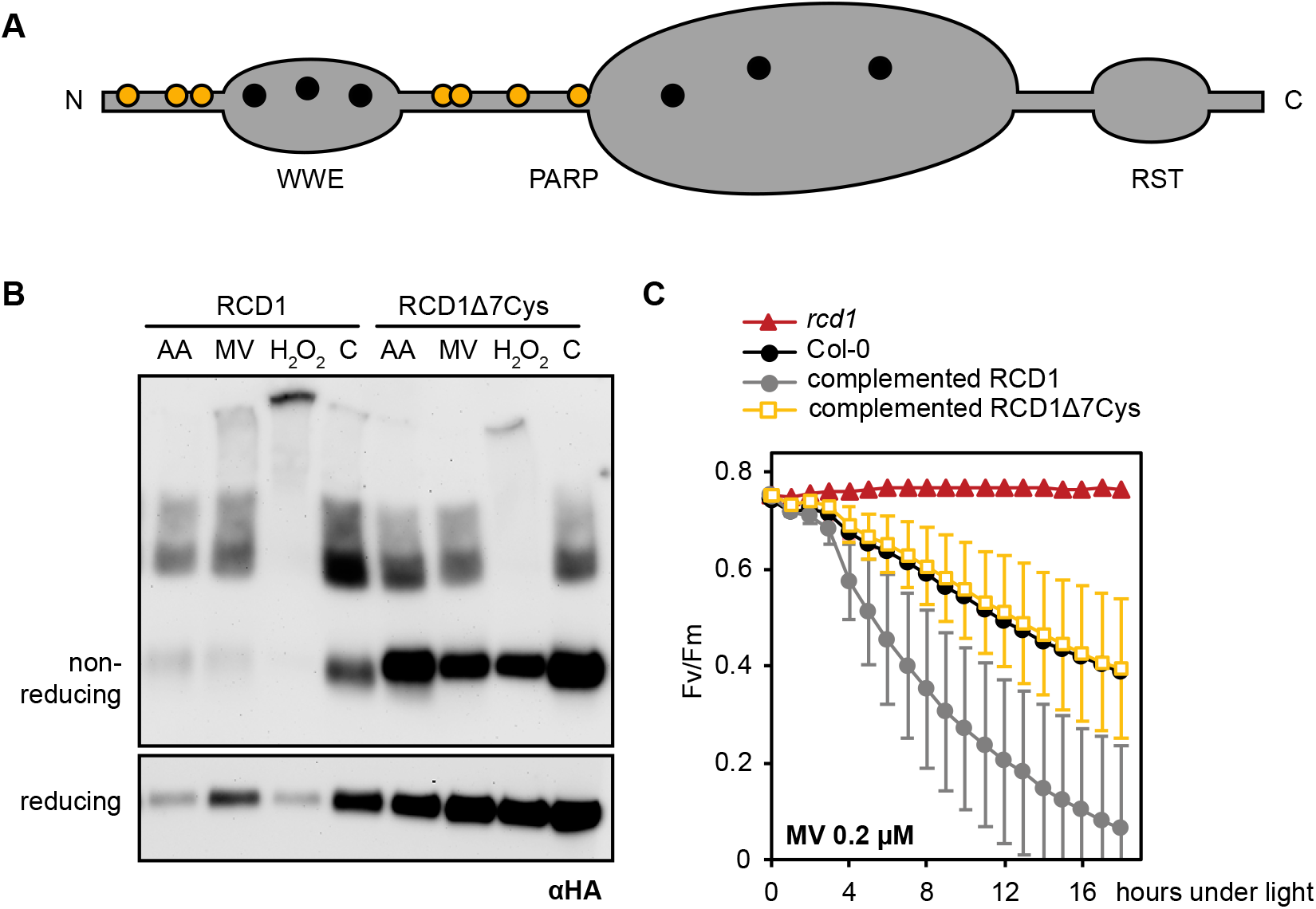
Characterization of the RCD1Δ7Cys mutant. (A) Domain structure of RCD1 with the positions of cysteine residues. Interdomain cysteines mutated in the RCD1Δ7Cys mutant are shown in yellow. (B) Formation of disulfide-bonded RCD1 complexes in the HA-tagged wild type RCD1 or the RCD1Δ7Cys line under chemical treatments *versus* water control (c). (C) MV-induced PSII photoinhibition in the RCD1Δ7Cys complemented line (mean ± SD, no significant difference observed between the complemented RCD1Δ7Cys line and Col-0, for full statistical analysis see Supplementary Table 5).

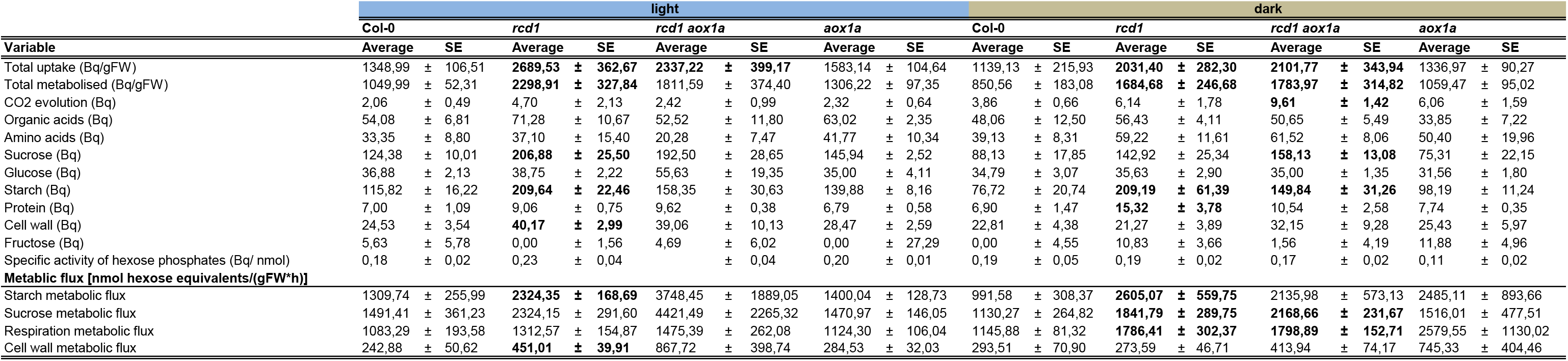

| Variable | light |  |  |  |  |  |  |  | dark |  |  |  |  |  |  |  |
| --- | --- | --- | --- | --- | --- | --- | --- | --- | --- | --- | --- | --- | --- | --- | --- | --- |
|  | Col-0 |  | *rcd1* |  | *rcd1 aox1a* |  | *aox1a* |  | Col-0 |  | *rcd1* |  | *rcd1 aox1a* |  | *aox1a* |  |
|  | Average | SE | Average | SE | Average | SE | Average | SE | Average | SE | Average | SE | Average | SE | Average | SE |
| Total uptake (Bq/gFW) | 1348,99 | ± 106,51 | **2689,53** | ± **362,67** | **2337,22** | ± **399,17** | 1583,14 | ± 104,64 | 1139,13 | ± 215,93 | **2031,40** | ± **282,30** | **2101,77** | ± **343,94** | 1336,97 | ± 90,27 |
| Total metabolised (Bq/gFW) | 1049,99 | ± 52,31 | **2298,91** | ± **327,84** | 1811,59 | ± 374,40 | 1306,22 | ± 97,35 | 850,56 | ± 183,08 | **1684,68** | ± **246,68** | **1783,97** | ± **314,82** | 1059,47 | ± 95,02 |
| CO2 evolution (Bq) | 2,06 | ± 0,49 | 4,70 | ± 2,13 | 2,42 | ± 0,99 | 2,32 | ± 0,64 | 3,86 | ± 0,66 | 6,14 | ± 1,78 | **9,61** | ± **1,42** | 6,06 | ± 1,59 |
| Organic acids (Bq) | 54,08 | ± 6,81 | 71,28 | ± 10,67 | 52,52 | ± 11,80 | 63,02 | ± 2,35 | 48,06 | ± 12,50 | 56,43 | ± 4,11 | 50,65 | ± 5,49 | 33,85 | ± 7,22 |
| Amino acids (Bq) | 33,35 | ± 8,80 | 37,10 | ± 15,40 | 20,28 | ± 7,47 | 41,77 | ± 10,34 | 39,13 | ± 8,31 | 59,22 | ± 11,61 | 61,52 | ± 8,06 | 50,40 | ± 19,96 |
| Sucrose (Bq) | 124,38 | ± 10,01 | **206,88** | ± **25,50** | 192,50 | ± 28,65 | 145,94 | ± 2,52 | 88,13 | ± 17,85 | 142,92 | ± 25,34 | **158,13** | ± **13,08** | 75,31 | ± 22,15 |
| Glucose (Bq) | 36,88 | ± 2,13 | 38,75 | ± 2,22 | 55,63 | ± 19,35 | 35,00 | ± 4,11 | 34,79 | ± 3,07 | 35,63 | ± 2,90 | 35,00 | ± 1,35 | 31,56 | ± 1,80 |
| Starch (Bq) | 115,82 | ± 16,22 | **209,64** | ± **22,46** | 158,35 | ± 30,63 | 139,88 | ± 8,16 | 76,72 | ± 20,74 | **209,19** | ± **61,39** | **149,84** | ± **31,26** | 98,19 | ± 11,24 |
| Protein (Bq) | 7,00 | ± 1,09 | 9,06 | ± 0,75 | 9,62 | ± 0,38 | 6,79 | ± 0,58 | 6,90 | ± 1,47 | **15,32** | ± **3,78** | 10,54 | ± 2,58 | 7,74 | ± 0,35 |
| Cell wall (Bq) | 24,53 | ± 3,54 | **40,17** | ± **2,99** | 39,06 | ± 10,13 | 28,47 | ± 2,59 | 22,81 | ± 4,38 | 21,27 | ± 3,89 | 32,15 | ± 9,28 | 25,43 | ± 5,97 |
| Fructose (Bq) | 5,63 | ± 5,78 | 0,00 | ± 1,56 | 4,69 | ± 6,02 | 0,00 | ± 27,29 | 0,00 | ± 4,55 | 10,83 | ± 3,66 | 1,56 | ± 4,19 | 11,88 | ± 4,96 |
| Specific activity of hexose phosphates (Bq/ nmol) | 0,18 | ± 0,02 | 0,23 | ± 0,04 |  | ± 0,04 | 0,20 | ± 0,01 | 0,19 | ± 0,05 | 0,19 | ± 0,02 | 0,17 | ± 0,02 | 0,11 | ± 0,02 |
| **Metabolic flux [nmol hexose equivalents/(gFW\*h)]** |  |  |  |  |  |  |  |  |  |  |  |  |  |  |  |  |
| Starch metabolic flux | 1309,74 | ± 255,99 | **2324,35** | ± **168,69** | 3748,45 | ± 1889,05 | 1400,04 | ± 128,73 | 991,58 | ± 308,37 | **2605,07** | ± **559,75** | 2135,98 | ± 573,13 | 2485,11 | ± 893,66 |
| Sucrose metabolic flux | 1491,41 | ± 361,23 | 2324,15 | ± 291,60 | 4421,49 | ± 2265,32 | 1470,97 | ± 146,05 | 1130,27 | ± 264,82 | **1841,79** | ± **289,75** | **2168,66** | ± **231,67** | 1516,01 | ± 477,51 |
| Respiration metabolic flux | 1083,29 | ± 193,58 | 1312,57 | ± 154,87 | 1475,39 | ± 262,08 | 1124,30 | ± 106,04 | 1145,88 | ± 81,32 | **1786,41** | ± **302,37** | **1798,89** | ± **152,71** | 2579,55 | ± 1130,02 |
| Cell wall metabolic flux | 242,88 | ± 50,62 | **451,01** | ± **39,91** | 867,72 | ± 398,74 | 284,53 | ± 32,03 | 293,51 | ± 70,90 | 273,59 | ± 46,71 | 413,94 | ± 74,17 | 745,33 | ± 404,46 |

**Supplementary Table 4.**
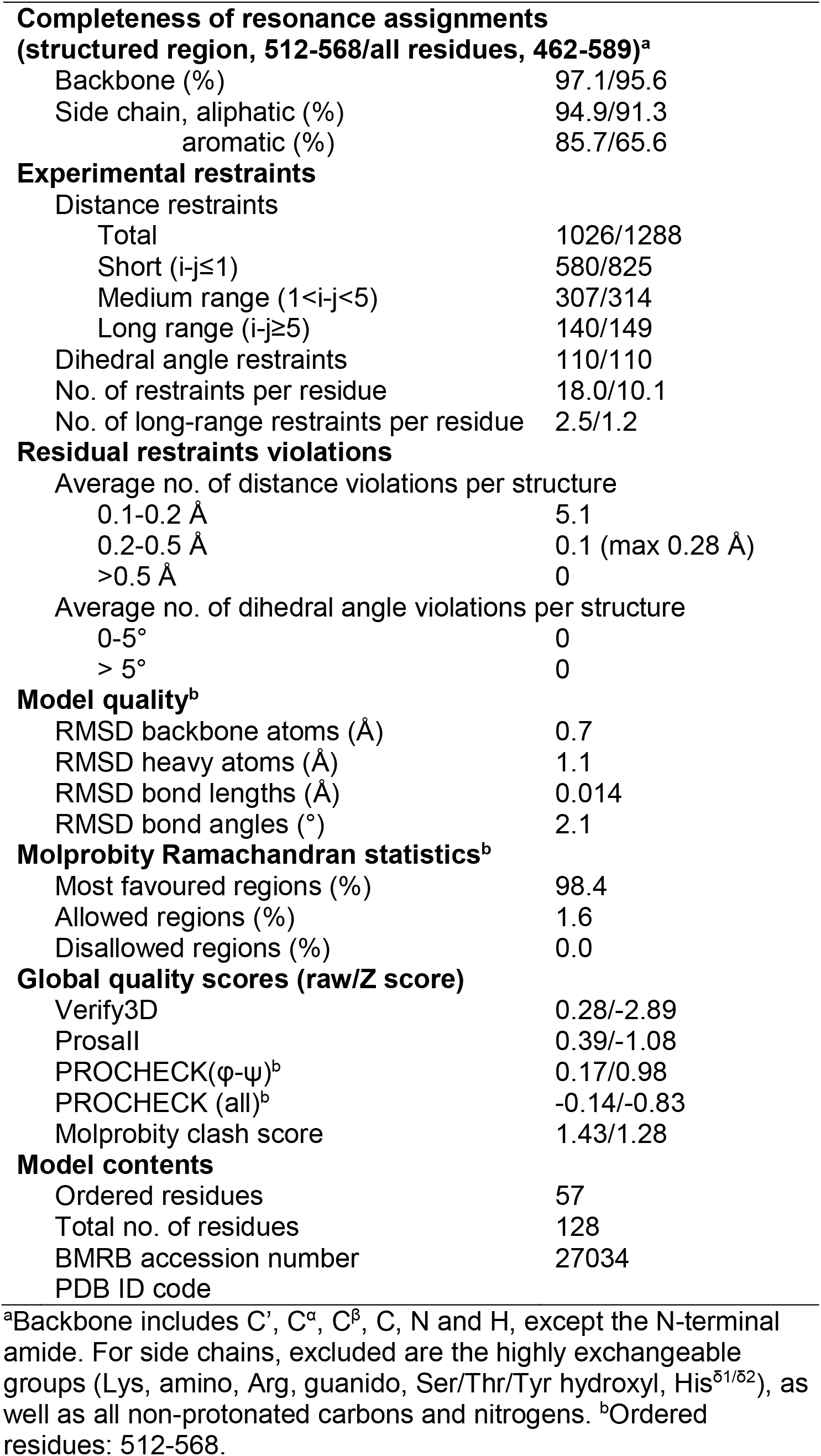
NMR constraints and structural statistics for the ensemble of the 15 lowest-energy structures of RCD1 RST.

**(structured region, 512-568/all residues, 462-589)<sup>a</sup>**
|  |  |
| --- | --- |
| Backbone (%) | 97.1/95.6 |
| Side chain, aliphatic (%) | 94.9/91.3 |
| aromatic (%) | 85.7/65.6 |
| **Experimental restraints** |  |
| Distance restraints |  |
| Total | 1026/1288 |
| Short ($i-j \leq 1$) | 580/825 |
| Medium range ($1 < i-j < 5$) | 307/314 |
| Long range ($i-j \geq 5$) | 140/149 |
| Dihedral angle restraints | 110/110 |
| No. of restraints per residue | 18.0/10.1 |
| No. of long-range restraints per residue | 2.5/1.2 |
| **Residual restraints violations** |  |
| Average no. of distance violations per structure |  |
| 0.1-0.2 Å | 5.1 |
| 0.2-0.5 Å | 0.1 (max 0.28 Å) |
| >0.5 Å | 0 |
| Average no. of dihedral angle violations per structure |  |
| 0-5° | 0 |
| > 5° | 0 |
| **Model quality<sup>b</sup>** |  |
| RMSD backbone atoms (Å) | 0.7 |
| RMSD heavy atoms (Å) | 1.1 |
| RMSD bond lengths (Å) | 0.014 |
| RMSD bond angles (°) | 2.1 |
| **Molprobit Ramachandran statistics<sup>b</sup>** |  |
| Most favoured regions (%) | 98.4 |
| Allowed regions (%) | 1.6 |
| Disallowed regions (%) | 0.0 |
| **Global quality scores (raw/Z score)** |  |
| Verify3D | 0.28/-2.89 |
| ProsaII | 0.39/-1.08 |
| PROCHECK(ϕ-ψ)<sup>b</sup> | 0.17/0.98 |
| PROCHECK (all)<sup>b</sup> | -0.14/-0.83 |
| Molprobit clash score | 1.43/1.28 |
| **Model contents** |  |
| Ordered residues | 57 |
| Total no. of residues | 128 |
| BMRB accession number | 27034 |
| PDB ID code |  |

<sup>a</sup>Backbone includes C', C<sup>α</sup>, C<sup>β</sup>, C, N and H, except the N-terminal amide. For side chains, excluded are the highly exchangeable groups (Lys, amino, Arg, guanido, Ser/Thr/Tyr hydroxyl, His<sup>δ1/δ2</sup>), as well as all non-protonated carbons and nitrogens. <sup>b</sup>Ordered residues: 512-568.

